# Sperm development requires TMC5-dependent mechanoresponsive calcium entry

**DOI:** 10.64898/2026.09.26.754653

**Authors:** W. Sharon Zheng, Lorena Roa-de la Cruz, John Duncan, Liam Slocomb, Betta Chopard, Michelina Kierzek, Fan Lin, Alexander Webb, Erin Jeffery, Celia M. Santi, Jung-Bum Shin, Sameer S. Bajikar, Gloria Shenynkman, Bimal N. Desai, Brian P. Hermann, Christopher B. Geyer, Seham Ebrahim

**Affiliations:** Center for Membrane and Cell Physiology, University of Virginia School of Medicine, Charlottesville, VA 22903, USA; Department of Molecular Physiology and Biological Physics, University of Virginia School of Medicine, Charlottesville, VA 22903, USA; Department of Neuroscience, Developmental and Regenerative Biology, The University of Texas at San Antonio, San Antonio, TX 78249, USA; Department of Pharmacology, University of Virginia School of Medicine, Charlottesville, VA 22903, USA; Department of Anatomy & Cell Biology, Brody School of Medicine at East Carolina University, Greenville, NC 27834, USA; Department of Obstetrics and Gynecology, Washington University in St. Louis, St. Louis, MO 63110, USA; Department of Neuroscience, University of Virginia School of Medicine, Charlottesville, VA 22903, USA; Depertment of Cell Biology, University of Virginia School of Medicine, Charlottesville, VA 22903, USA

## Abstract

Spermiogenesis, the transformation of round spermatids into streamlined, fertilization-competent sperm, represents cellular morphogenesis at its most extreme^1,2^. Despite decades of effort, spermiogenesis remains difficult to recapitulate outside the testis^2,3^, highlighting the importance of cues within the native tissue environment. The nature of these cues, however, remains poorly understood. Here, we discover that mechanical stimulation elicits Ca^2+^ entry in spermatids and identify transmembrane channel-like protein 5 (TMC5) as a principal mediator of this response. In both mouse and human testes, TMC5 is selectively expressed in spermatids and localized to the plasma membrane. TMC5 associates with TMC7 and calcium- and integrin-binding (CIB) proteins, mirroring TMC–CIB mechanotransduction complexes in sensory organs^4,5^. Mechanistically, loss of *Tmc5* blocks mechanically evoked Ca^2+^ entry, leading to disruptions in cytoskeletal remodeling and failure of CREMτ-dependent transcriptional programs required for spermiogenesis^6^. Thus, TMC5 is required to couple mechanical cues to sperm development, with direct implications for male infertility.

## Introduction

Cell differentiation is often described as a cell-intrinsic program, yet many developmental transitions depend on instructive signals from specialized tissue microenvironments. Spermiogenesis-the final phase of sperm development-provides a striking example. Within the seminiferous epithelium, haploid round spermatids undergo extensive membrane and cytoskeletal remodeling, acrosome and flagellum formation, nuclear compaction, and substantial cytoplasmic elimination to become fertilization-competent spermatozoa^1^. Although numerous transcriptional regulators and structural components required for this dramatic transformation have been identified, the morphogenetic program of spermiogenesis has not been fully recapitulated outside the testis^2,3^. This reveals that the seminiferous epithelium provides developmental cues essential for spermiogenesis. However, the identities of these cues, and the mechanisms by which spermatids translate them into the cellular remodeling events that drive spermiogenesis, remain largely unknown.

Spermatids develop within a mechanically dynamic environment, and their development is particularly vulnerable to disruptions in calcium (Ca²⁺) homeostasis^7–9^. Spermatids remain closely associated with specialized somatic supporting Sertoli cells as they move through the adluminal compartment of the seminiferous epithelium, while also undergoing profound changes in cell shape, volume, surface area, and membrane organization^1^. These interactions with the surrounding seminiferous epithelium^10^ likely expose spermatids to mechanical forces that are expected to generate fluctuations in plasma-membrane tension, although neither premise has been tested. In several other cell types, changes in membrane tension or deformation are converted into Ca^2+^ signals by mechanically activated ion channels, which ultimately regulate cytoskeletal organization, gene expression, and cell fate^11,12^. Whether developing spermatids sense mechanical cues, and whether such mechanotransduction contributes to their differentiation, has remained unknown.

The molecular identity of a potential spermatid mechanotransduction apparatus is similarly unclear. Surprisingly, canonical mechanically activated channels, including PIEZO and TRPV family members, are minimally expressed or undetectable in post-meiotic spermatids^13^. However, members of the transmembrane channel-like (TMC) family of putative mechanosensitive ion channels, specifically *Tmc5* and *Tmc7,* are selectively expressed in developing spermatids in both mouse and human testis^13^. The TMC family comprises eight conserved multipass membrane proteins, of which TMC1 and TMC2 form core components of the mechanotransduction channel in inner-ear hair cells^14,15^. Together with calcium- and integrin-binding (CIB) proteins, these channels convert stereociliary deflection into ion influx^4,14^. The remaining mammalian TMC family proteins were also recently proposed to be mechanically gated ion channels^16^. TMC5 shares structural features with TMC1 within the putative ion-conducting cavity, and molecular-dynamics simulations suggest that it can support calcium permeation^17^. TMC7 has independently been linked to male fertility, although it has been proposed to function intracellularly in Golgi homeostasis and acrosome biogenesis^18,19^. Whether TMC proteins form a mechanosensitive complex at the spermatid plasma membrane has not been established.

Here, we show that mechanical stimulation elicits Ca^2+^ entry in developing spermatids and identify TMC5, in complex with TMC7 and CIB proteins at the spermatid plasma membrane, as the principal mediator of this mechanotransduction. Loss of TMC5 attenuates Ca^2+^ entry, disrupting calcium/calmodulin-dependent cytoskeletal remodeling and the CREMτ-dependent transcriptional program required for spermatid elongation, and results in complete male infertility characterized by non-obstructive azoospermia (NOA). Because the genetic basis of NOA remains unresolved in the majority of patients^20^, and TMC5 expression is conserved in human spermatids, our findings also nominate TMC5 as a candidate for future evaluation in human NOA.

## Results

### Developing spermatids exhibit PIEZO1-independent mechanosensitive Ca²⁺ entry

Spermiogenesis entails extensive changes in cell shape, volume, surface area, and membrane organization (Fig. 1a), transformations that are expected to generate fluctuations in plasma-membrane tension^21^. We therefore asked whether developing spermatids respond directly to mechanical stimulation and used hypotonic swelling as a non-contact, spatially distributed mechanical stimulus^22–24^. Isotonic and hypotonic buffers were matched in ionic composition, with osmolarity reduced from approximately 314 to 214 mOsm by omitting mannitol during hypotonic stimulation (Fig. 1b). This design separated the effects of osmotic swelling from those of altered extracellular ion concentrations. Intracellular Ca^2+^ was monitored using Fura-2, a ratiometric indicator whose excitation shifts upon Ca^2+^ binding. An increase in the F_340_/_F380_ ratio therefore indicates an increase in intracellular Ca^2+^.

**Figure 1.**
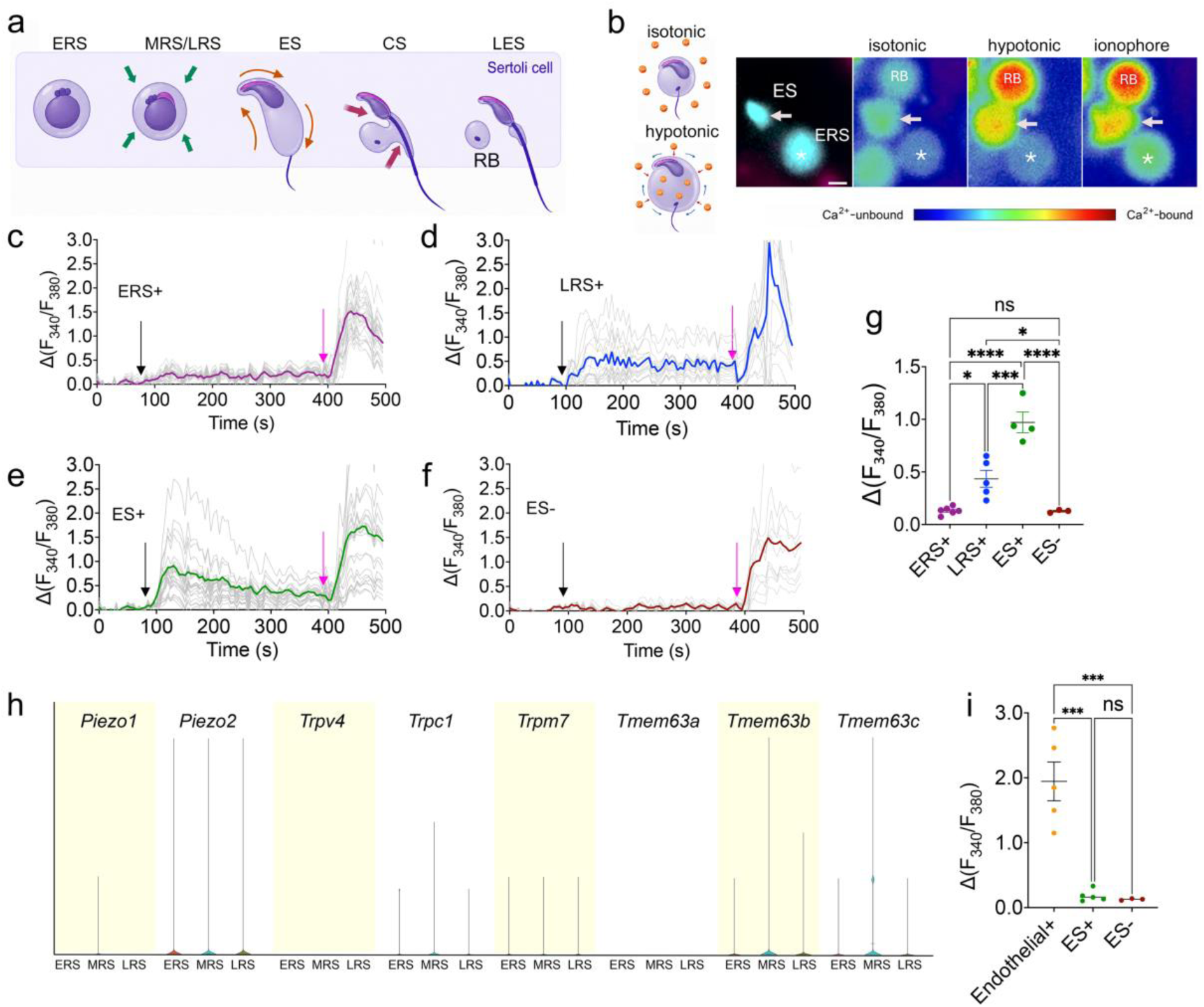
Developing spermatids exhibit mechanosensitive Ca²⁺ influx. **a,** The steps of spermatid development occur in close juxtaposition with Sertoli cell. Arrows indicate likely membrane tension and remodeling forces. ERS, early round (step 1-3); MRS/LRS, mid/late round (step 4-6/7-8); ES, elongating (step 9-11); CS, condensing spermatid (step 12-14); LES, late elongated spermatid (step 15-16); RB, residual body. **b,** Hypotonic swelling schematic and representative images identifying an ES (arrow), ERS (asterisk) and residual body (RB). Scale bar, 5 μm. **c–f,** Fura-2 traces from ERS⁺ (c), LRS⁺ (d), ES⁺ (e) and ES⁻ (f) cells during hypotonic stimulation (black arrow) and ionophore addition (pink arrow). Gray, individual traces; color, mean. **g,** Peak hypotonicity-evoked responses. **h,** Expression of canonical mechanosensitive channels across ERS, MRS and LRS populations. **i,** Peak Yoda1-evoked responses in endothelial, ES⁺ and ES⁻ cells. Superscript + indicates Ca²⁺-containing buffer; superscript − indicates Ca²⁺-free buffer. Dots represent independent experiments. ns, not significant; *P < 0.05, **P < 0.01, ***P < 0.001 and ****P < 0.0001.

Early round spermatids (ERS), late round spermatids (LRS) and elongating spermatids (ES) showed stable baseline F_340_/_F380_ in isotonic buffer before stimulation (Fig. 1c-f). Hypotonic stimulation (black arrow, Fig. 1c-f) elicited a robust increase in intracellular Ca²⁺ in both LRS and ES (Fig. 1d,e). The response amplitude was greater in ES than in LRS (Fig. 1g). Conversely, no Ca^2+^ response to hypotonic stimulation was detected in ERS (Fig. 1c,g). Together, these data reveal that Ca^2+^ influx in response to mechanical stimuli is acquired as spermatids mature, emerging just before elongation and increases as spermiogenesis proceeds (Fig. 1a,g).

Swelling-evoked Ca^2+^ responses were abolished in Ca^2+^-free extracellular buffer (Fig. 1f,g), demonstrating that they require Ca^2+^ entry across the plasma membrane. Ionophore nevertheless increased cytoplasmic Ca^2+^ under Ca^2+^-free conditions, confirming spermatid viability and the presence of releasable intracellular Ca^2+^ pools. Thus, the absence of mechanically evoked responses in Ca^2+^-free buffer cannot be attributed to loss of cell viability or depletion of intracellular Ca^2+^ stores.

We next asked whether this response was mediated by a canonical mechanically activated ion channel. Transcriptomic analysis across post-meiotic spermatid populations revealed little to no detectable expression of canonical mechanosensitive channels, including *Piezo1*, *Piezo2*, *Trpv4*, *Trpc1*, *Trpm7* or *Tmem63* in ERS, MRS, or LRS (Fig. 1h, Extended Data Fig. 1). We tested PIEZO1 function directly using Yoda1, a selective PIEZO1 agonist^25^. Yoda1 elicited a robust increase in F_340_/_F380_ ratio in endothelial cells used as a positive control but produced no detectable response in ESs maintained in Ca^2+^-containing buffer (Fig. 1i). The ES response remained indistinguishable from that measured in Ca^2+^-free conditions.

These gene expression and functional data argue that mechanically evoked Ca²⁺ entry in developing spermatids is independent of PIEZO1. These results identify mechanical force as a previously unrecognized input into spermatid development.

#### TMC5 and TMC7 are selectively expressed in spermatids and localize to the plasma membrane

In contrast to the minimal or undetectable expression of canonical mechanically activated ion channels, both *Tmc5* and *Tmc7* genes were selectively and robustly expressed in postmeiotic spermatids at the developmental stages when mechanically evoked Ca²⁺ influx was observed. Both transcripts were minimally detected in preceding mitotic spermatogonia and meiotic spermatocytes (Fig. 2a,c) ^13^. Importantly, this spermatid-enriched expression pattern was also conserved in humans (Fig. 2b,d).

**Figure 2.**
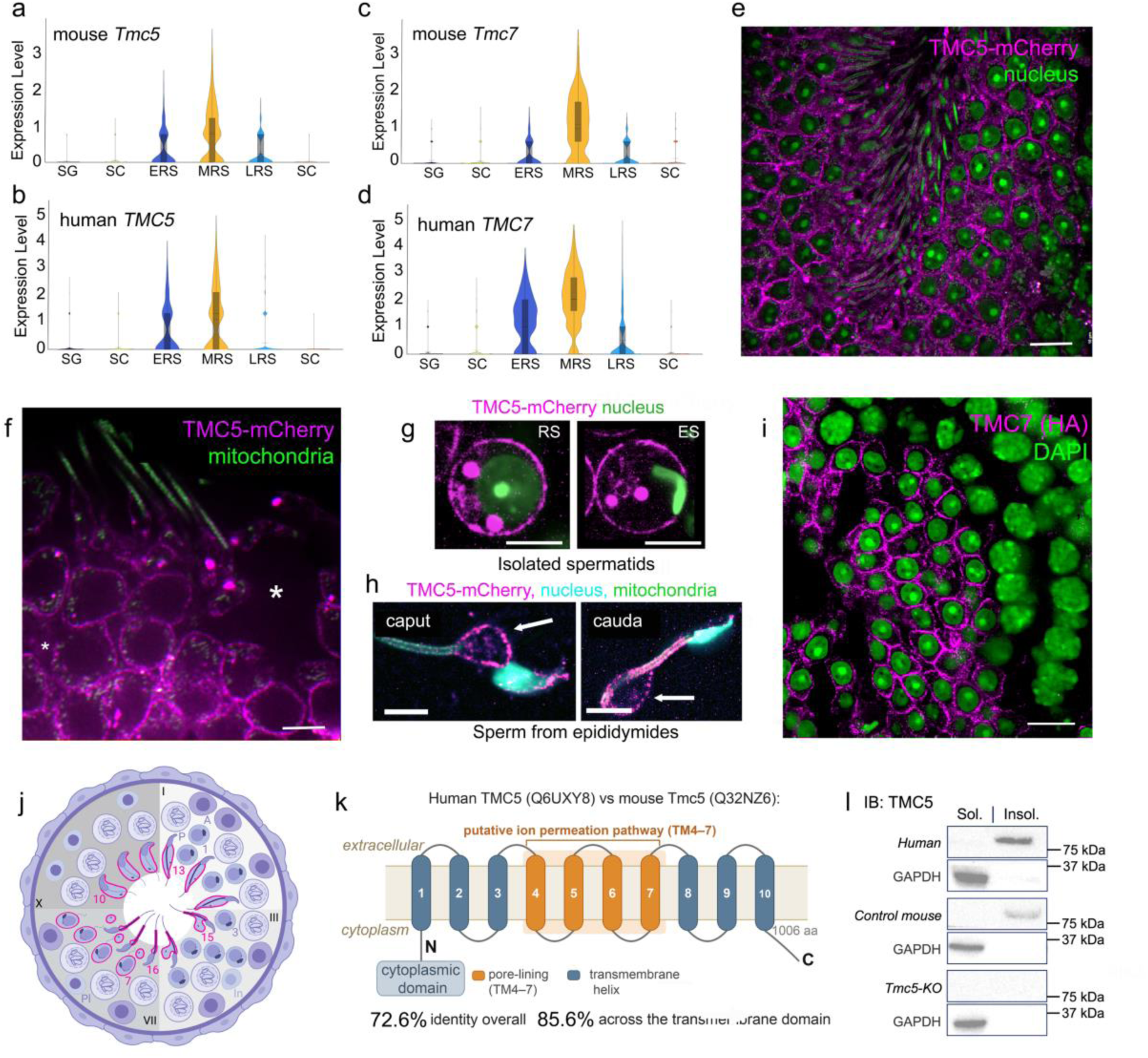
TMC5 is selectively expressed in developing spermatids and localizes to the plasma membrane. **a–d**, *Tmc5*/*TMC5* and *Tmc7*/*TMC7* expression across mouse (a,c) and human (b,d) testicular cell populations. **e-f,***Tmc5*–mCherry seminiferous-tubule sections showing TMC5–mCherry (magenta) with nuclei (e) or mitochondria (f) in green. Asterisks indicate Sertoli cells. **g**, TMC5–mCherry localization in isolated round (RS) and elongating (ES) spermatids. Nuclei, green. **h**, TMC5–mCherry localization in caput and cauda epididymal sperm. Nuclei, cyan; mitochondria, green. Arrows indicate cytoplasmic droplets. **i**, TMC7–HA localization in a seminiferous tubule. TMC7–HA, magenta; DAPI, green. **j**, Schematic of TMC5 and TMC7 (magenta) expression during spermatid development. **k**, Predicted TMC5 topology showing ten transmembrane helices and the putative TM4–TM7 ion-permeation pathway. Mouse and human TMC5 share 72.6% overall sequence identity and 85.6% identity across the transmembrane domain. **l**, TMC5 immunoblot of soluble (Sol.) and insoluble (Insol.) fractions from human, mouse and *Tmc5*-KO testes. Scale bars, 5 μm, except i, 10 μm

We next examined the subcellular localization of TMC5 and TMC7 proteins in developing spermatids. Because antibodies against TMC5 and TMC7 did not work well for immunofluorescence staining, we used a *Tmc5*–mCherry knock-in reporter mouse^17^ and generated a *Tmc7*–HA reporter mouse line (Extended Data Fig. 2a). In confocal images of seminiferous tubule cross-sections, we readily detected TMC5–mCherry signal continuously from MRS through LES stages (Fig. 2e,f and Extended Data Fig. 2b,c), spanning the developmental window in which mechanosensitive Ca²⁺ responses were observed (Fig. 1). As predicted from the transcriptome data, TMC5–mCherry was undetectable in precursor spermatogonia and spermatocytes (Extended Data Fig. 2b) as well as in somatic Sertoli cells (Fig. 2f, asterisk). In isolated spermatids, TMC5–mCherry was enriched at the plasma membrane of both round and elongating spermatids (Fig. 2g). TMC5–mCherry was also detected on epididymal spermatozoa, from both caput and cauda epididymides, and enriched at the cytoplasmic droplet (Fig. 2h), revealing that TMC5 is retained on sperm during epididymal transit. Although TMC7 was previously proposed to function at the Golgi, endogenous TMC7-HA localized predominantly to the spermatid periphery, similar to TMC5 and consistent with plasma-membrane localization (Fig. 2i). The developmental stages across which TMC5 and TMC7 are expressed and localized to the spermatid periphery are summarized in Fig. 2j. We focused the remainder of the study on TMC5 because its function during sperm development is unexplored.

Importantly, mouse and human TMC5 share high sequence and structural similarity across the full protein length (Fig. 2k and Extended Data Fig. 2d), supporting the translation of results from genetic mouse models to humans. We validated TMC5 protein expression biochemically by immunoblot of testis lysate fractions (Fig. 2l). Endogenous TMC5 was enriched in the insoluble fraction from both mouse and human testes, consistent with its predicted integral membrane topology, and was absent from *Tmc5*-knockout (KO) testis lysates, confirming antibody specificity.

Together, these transcriptomic, biochemical, and imaging data establish TMC5 as an evolutionarily conserved spermatid-enriched integral membrane protein that localizes to the plasma membrane throughout the spermiogenesis developmental window of mechanosensitive calcium signaling, consistent with a direct role in mechanically evoked calcium entry.

#### Loss of TMC5 blocks mechanoresponsive calcium influx and spermatid elongation

To define the functional requirement for TMC5 in spermiogenesis, we generated *Tmc5*-KO mice by CRISPR–Cas9-mediated introduction of a 44-bp deletion within exon 8 (Fig. 3a). The loss of TMC5 protein was verified by immunoblotting (Fig. 2l). *Tmc5*-KO males had significantly smaller testes relative to body weight (Fig. 3b). In contrast to controls, *Tmc5*-KO males produced no offspring, demonstrating complete infertility (Fig. 3c). Histological analyses of PAS-stained sections revealed that control testes were grossly normal, and epididymides contained abundant epididymal sperm (Fig. 3d,e). Conversely, *Tmc5*-KO testes largely lacked ES, CS, or LES and epididymides lacked sperm entirely (Fig. 3d,e).

**Figure 3.**
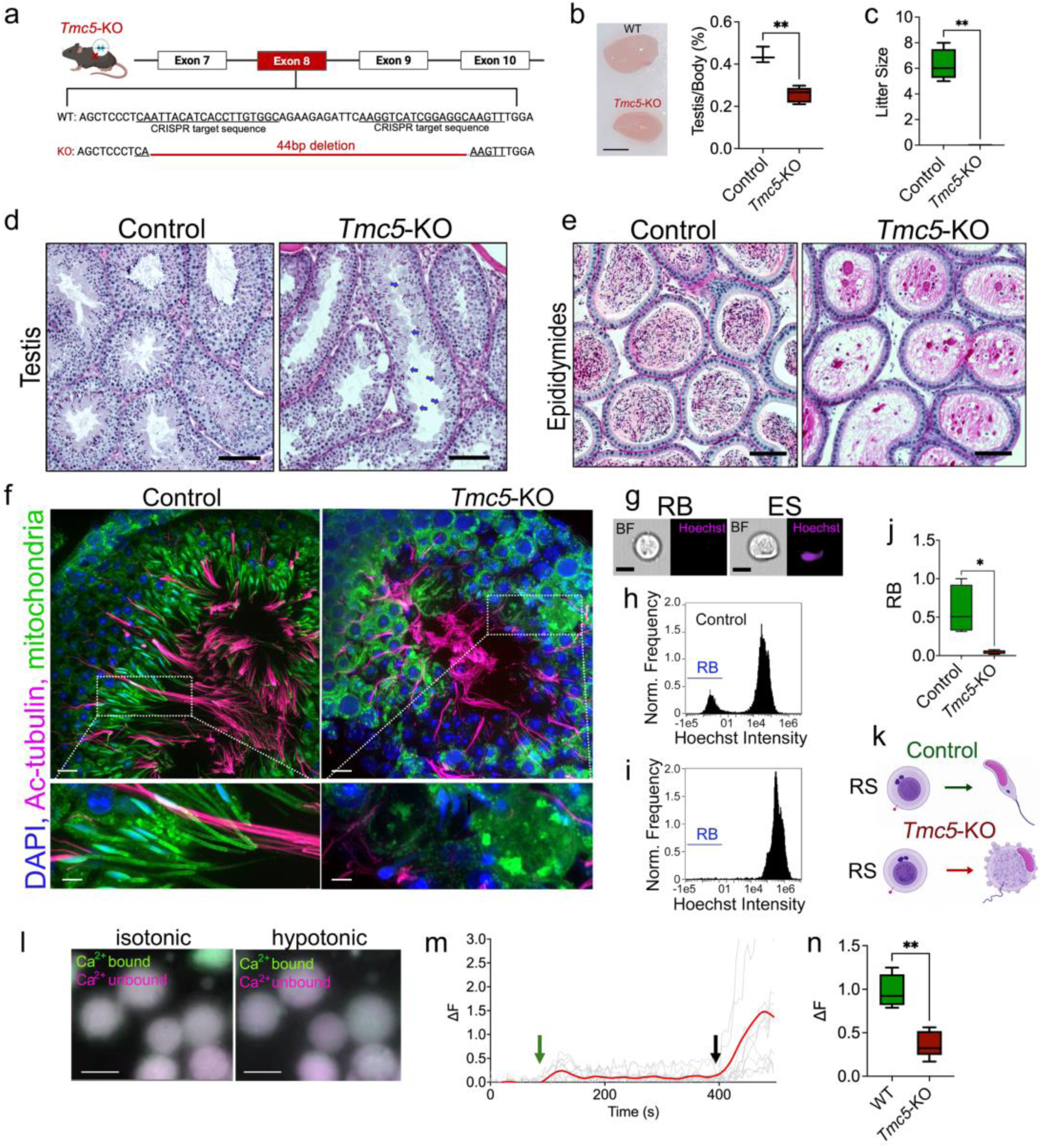
TMC5 is required for mechanically evoked calcium influx and spermatid elongation. **a,** CRISPR-Cas9 strategy used to generate *Tmc5*-KO mice. Exon 8 was targeted with two guide RNAs (underlined), producing a 44-bp deletion and frameshift. **b,** Representative testes and testis-to-body weight ratios from control and *Tmc5*-KO males. Scale bar, 5 mm. **c,** Litter sizes from control and *Tmc5*-KO males crossed with control females. *Tmc5*-KO males produced no offspring. **d,** PAS-stained testis sections from control and *Tmc5*-KO mice. Seminiferous tubule stages are indicated by Roman numerals. Arrows indicate tubules depleted of elongating spermatids. Scale bars, 50 μm. **e,** Histological sections of cauda epididymides from control and *Tmc5*-KO mice. Control tubules contain abundant mature sperm, whereas *Tmc5*-KO tubules lack sperm. Scale bars, 50 μm. **f,** Confocal images of control and *Tmc5*-KO seminiferous tubules stained with DAPI (blue), acetylated α-tubulin (magenta), and a mitochondrial marker (green). Lower panels show higher-magnification views of the boxed regions. Scale bars, 10 μm; magnified views, 5 μm. **g,** Representative brightfield (BF) and Hoechst images acquired by imaging flow cytometry showing a residual body (RB) and an elongating spermatid (ES), illustrating the morphological and fluorescence criteria used to identify the anucleate RB population. Scale bars, 5 μm. h,i, Hoechst-intensity distributions of germ-cell suspensions from control (h) and *Tmc5*-KO (**i**) testes. The low-Hoechst RB population (blue) is present in control testes but absent from *Tmc5*-KO testes. **j,** RB abundance in control and *Tmc5*-KO germ-cell suspensions. **k,** Schematic of spermatid development in control and *Tmc5*-KO mice. Control round spermatids (RS) elongate to form elongated spermatids, whereas *Tmc5*-KO round spermatids fail to elongate. **l, Representative Fura-2 images of Tmc5-KO ES in isotonic and hypotonic buffer. Ca²⁺-bound (340 nm), green; Ca²⁺-unbound (380 nm), magenta. Scale bars, 10 μm. m, Fura-2 traces (ΔF) from Tmc5-KO ES during hypotonic stimulation (green arrow) and ionophore addition (black arrow). Gray, individual traces; red, mean. n, Peak hypotonicity-evoked ΔF in control and Tmc5-KO ES.** * P < 0.05, **P <9

Confocal imaging of seminiferous tubules confirmed a striking disruption of spermatid development in *Tmc5*-KO testes compared to littermate controls (Fig. 3f). Whereas control tubules exhibited orderly progression through the spermiogenesis program with abundant round, elongating, and condensing spermatids (Fig. 3f), *Tmc5*-KO testes contained abundant round spermatids but few elongating and fewer condensing spermatids, revealing significant failure to either initiate or sustain elongation (Fig. 3f,k). Consistent with the elongation block, *Tmc5*-KO tubules showed a marked increase in TUNEL positive germ cells (Extended Data Fig. 3a,b) starting during the CS stage.

To quantify the extent of the spermatid elongation defect, we isolated germ cell suspensions from control and *Tmc5*-KO testes and assessed formation of residual bodies (extruded cytoplasm during normal spermiogenesis, Fig. 1a) by imaging flow cytometry. Residual bodies were distinguishable from nucleated elongating spermatids and identified as anucleate cytoplasmic bodies by their characteristic size and absence of Hoechst signal, distinguishable from nucleated elongating spermatids (Fig. 3g). In control preparations, a distinct residual body population was readily detected at low Hoechst intensity, whereas *Tmc5*-KO preparations showed near-complete absence of this population (Fig. 3h,i), indicating that the cytoplasmic extrusion step of late elongating spermatid (LES) fails to occur. The absence of residual bodies is consistent with spermatids failing to reach the LES stage at which cytoplasmic remodeling and shedding take place^1^ (Fig. 3j).

Finally, we asked whether TMC5 is required for mechanically evoked calcium signaling in KO spermatids. The robust Ca^2+^ influx induced by hypotonic swelling in control spermatids (Fig. 1e) was markedly diminished in *Tmc5*-KO spermatids (Fig. 3l–n) while still responsive to ionophore. Together, these findings identify TMC5 as a principal mediator of mechanically evoked Ca^2+^ entry in spermatids and link its loss to a complete block in spermatid elongation.

#### Loss of TMC5-dependent calcium signaling disrupts cytoskeletal dynamics during spermatid elongation

Because spermatogenesis proceeds asynchronously in adult testes, assigning reproductive phenotypes to specific developmental transitions can be challenging in knockout animals. To map the precise temporal onset of spermatid defects in *Tmc5*-KO mice, we synchronized spermatogenesis by manipulating retinoic acid (RA) signaling, as previously described (Fig. 4a)^26–28^. As expected, testes from mice with synchronized spermatogenesis contained a uniform cohort of spermatids at defined developmental steps, as confirmed by staining of cytoskeletal, nuclear and acrosomal markers (Fig. 4b,c).

**Figure 4.**
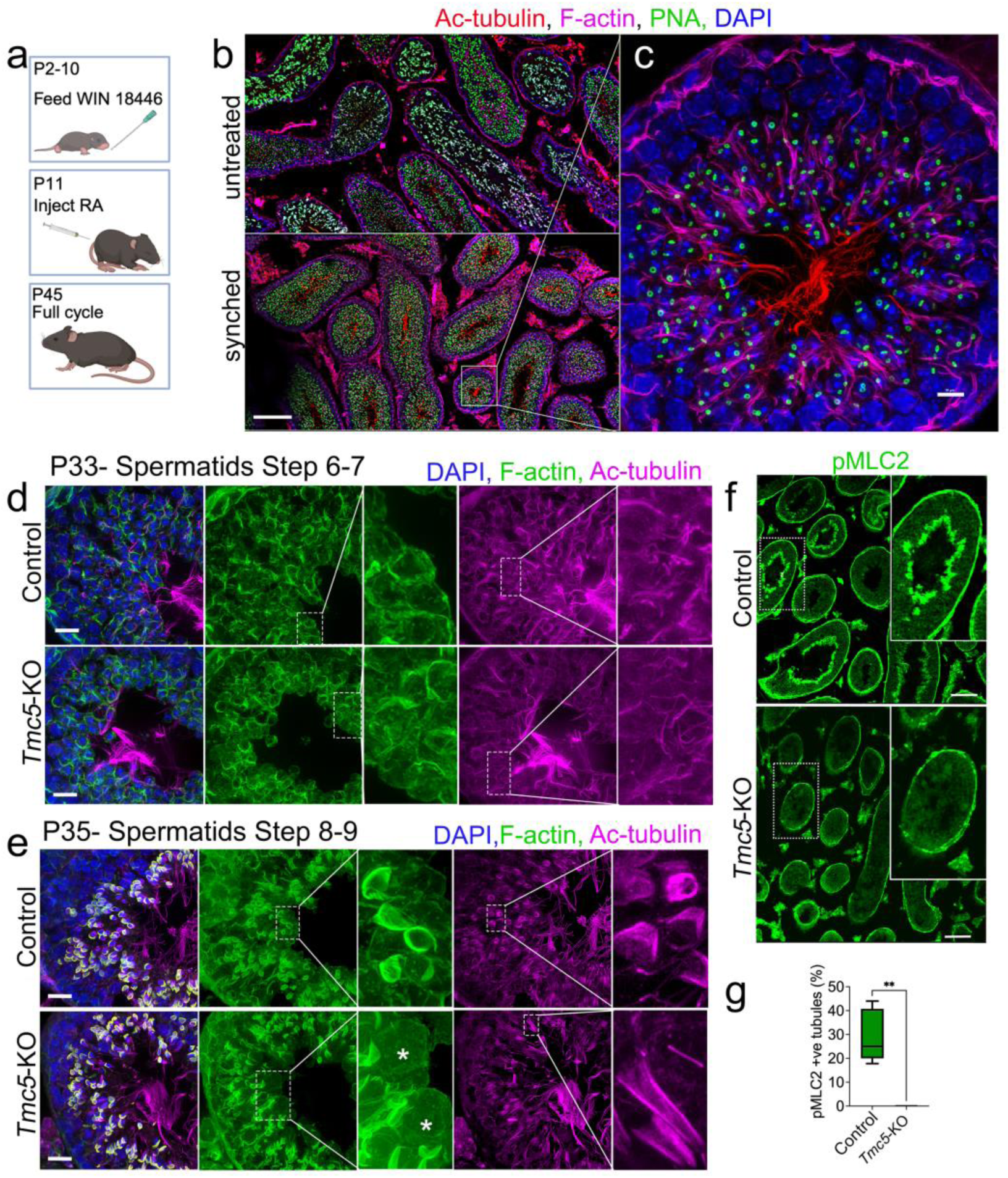
TMC5 is required for manchette organization and actomyosin activation in elongating spermatids. **a,** Schematic of the spermatogenesis synchronization protocol. Mice were fed WIN 18446 from P2–10 to block RA signaling, injected with exogenous RA at P11, and analyzed at subsequent defined timepoints. **b,** Confocal images of unsynchronized (untreated, top) and synchronized (bottom) adult seminiferous tubules stained for acetylated α-tubulin (red), F-actin (magenta), PNA-lectin (green), and DAPI (blue). Scale bar, 50 μm. **c,** High-magnification confocal image of a representative synchronized seminiferous tubule cross-section. Staining as in b. Scale bar, 10 μm. d, Confocal images of control (top) and *Tmc5*-KO (bottom) synchronized testes at P33 (spermatid steps 6–7) stained for DAPI (blue), F-actin (green), and Ac-tubulin (magenta). Control and *Tmc5*-KO spermatids are morphologically indistinguishable at this stage. Insets show higher-magnification views of boxed regions. Scale bar, 10 μm. **e,** Confocal images of control (top) and *Tmc5*-KO (bottom) synchronized testes at P35 (spermatid steps 8–9), stained as in d. Control spermatids display organized manchette structures and F-actin networks; *Tmc5*-KO spermatids show disorganized cytoskeletal architecture and failure to elongate. Asterisks indicate enlarged, undifferentiated cells. Insets show higher-magnification views of boxed regions. Scale bar, 10 μm. **f,** Confocal images of phosphorylated MLC2 (pMLC2; green) in control (top) and *Tmc5*-KO (bottom) adult seminiferous tubules. pMLC2 is enriched in the luminal region of control tubules, consistent with actomyosin activation in elongating spermatids; signal is absent across all tubule stages in *Tmc5*-KO testes. Insets show higher-magnification views of boxed regions. Scale bar, 50 μm. **g,** Percentage of pMLC2-positive tubules in control and *Tmc5*-KO testes. (n = 4

The dramatic morphological changes during spermiogenesis are driven by the remodeling of both microtubule and actin-based cytoskeletal structures. Specifically, cytoplasmic remodeling driven by actomyosin-contractility and nuclear shaping driven by a transient microtubule-structure called the manchette^27,29^. Because Ca^2+^/calmodulin signaling regulates manchette assembly^30^ and actomyosin contractility during cellular remodeling and tissue morphogenesis^31,32^, we next asked whether these processes require TMC5-dependent Ca^2+^ influx.

To address this question, we examined microtubule and actin organization in synchronized control and *Tmc5*-KO testes by immunofluorescence staining for acetylated tubulin and F-actin. Round spermatids until steps 6-7 from control and *Tmc5*-KO mice were morphologically indistinguishable (Fig. 4d), indicating mitotic proliferation of spermatogonia, meiosis of spermatocytes and formation of ERS proceed normally in the absence of TMC5. By steps 8-9 (LRS and ES, respectively), *Tmc5*-KO spermatids began to show defects in cytoskeletal organization (Fig. 4e), identifying the RS - ES transition as the major point of failure following TMC5 loss. Specifically, *Tmc5*-KO ESs had disorganized manchettes accompanied by impaired nuclear shaping and elongation (Fig. 4e). Phalloidin staining also revealed a markedly disorganized F-actin network in *Tmc5*-KO ESs, which appeared substantially enlarged relative to controls (Fig. 4e, insets), consistent with failed cytoplasmic elimination. Thus, the functional requirement for TMC5 emerges as spermatids initiate cytoskeletal transformations that drive elongation.

We next sought a molecular readout linking TMC5-dependent Ca^2+^ entry to actomyosin remodeling. Ca^2+^-bound calmodulin activates myosin light chain kinase (MLCK), which phosphorylates myosin regulatory light chain 2 (MLC2), promoting non-muscle myosin II assembly and contractility^33^. Phosphorylated MLC2 (pMLC2) therefore provides a proximal readout of the Ca^2+^/calmodulin–MLCK pathway through which TMC5-dependent Ca^2+^ influx could regulate actomyosin dynamics.

In control adult testes, pMLC2 was enriched in the luminal region of seminiferous tubules, where LES reside (Fig. 4f,g), coincident with the actomyosin-dependent cytoplasmic remodeling that accompanies cytoplasmic elimination and residual body formation. In contrast, pMLC2-positive spermatids were absent from *Tmc5*-KO testes (Fig. 4f,g), consistent with their arrest before activation of this terminal contractile program (Fig. 3f–j).

Together, these findings place the onset of the *Tmc5*-KO phenotype at the round-to-elongating spermatid transition and identify defects in both manchette organization and actomyosin remodeling. They support a model in which TMC5-dependent Ca^2+^ entry engages Ca^2+^/calmodulin-regulated cytoskeletal pathways required for spermatid elongation and cytoplasmic reorganization. These results prompted us to define the molecular complex through which TMC5 mediates mechanosensitive Ca^2+^ entry at the spermatid plasma membrane.

#### TMC5 associates with TMC7 and CIB proteins at the spermatid plasma membrane

TMC1- and TMC2-containing assemblies in auditory and vestibular hair cells^14,15,34^, operate in concert with accessory factors, including calcium- and integrin-binding (CIB) proteins, that support TMC channel localization, stability, and function^4,35,36^. We therefore reasoned that TMC5-dependent Ca^2+^ entry in spermatids similarly requires a specialized protein complex at the plasma membrane. Consistent with this idea, individual deletion of *Tmc5*, *Tmc7*, *Cib1*, or *Cib4* caused spermatid differentiation defects and male infertility, providing genetic support for the concept that the encoded proteins function in a shared complex^18,19,37,38^. *Cib1*/*CIB1* is expressed across male germ cells, while *Cib4*/*CIB4* shares the restricted spermatid-specific pattern of expression of *Tmc5*/*TMC5* and *Tmc7*/*TMC7*^13^ (Fig. 5a,b).

**Figure 5.**
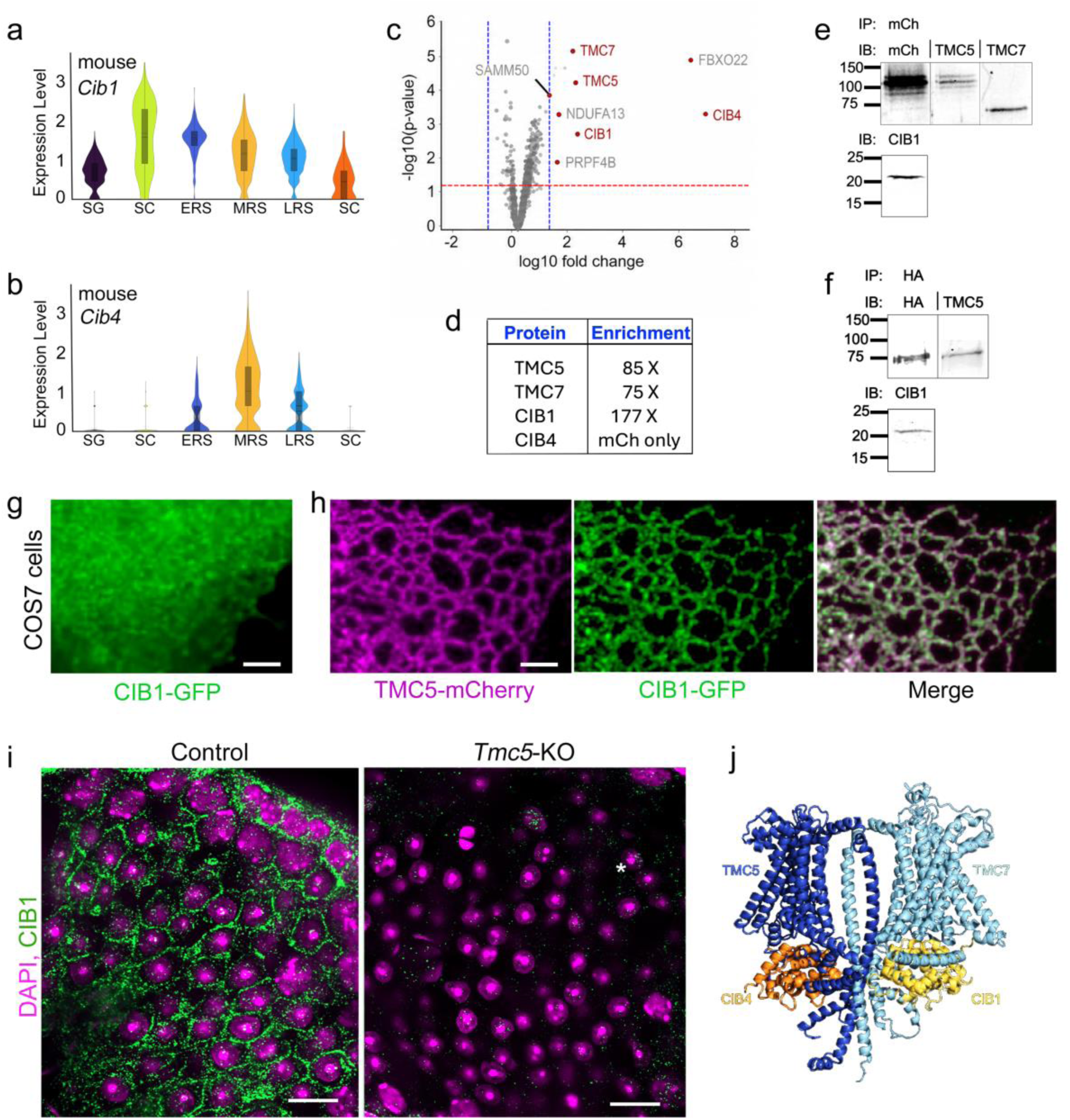
TMC5 assembles with TMC7 and CIB proteins at the spermatid plasma membrane. **a,b,** Cib1 (a) and Cib4 (b) expression across mouse testicular cell populations. **c**, Volcano plot of proteins enriched in TMC5–mCherry affinity purifications. Dashed lines indicate significance and fold-change thresholds. **d,** Enrichment of selected TMC5-associated proteins. **e,** TMC5–mCherry co-immunoprecipitation followed by immunoblotting for mCh, TMC5, TMC7 and CIB1. **f,** TMC7–HA co-immunoprecipitation followed by immunoblotting for HA, TMC5 and CIB1. **g**, CIB1–GFP expressed in COS7 cells is distributed diffusely throughout the cytoplasm. **h,** Co-expression of TMC5–mCherry and CIB1–GFP in COS7 cells. TMC5–mCherry is retained in the endoplasmic reticulum; CIB1–GFP redistributes to co-localize with TMC5–mCherry in the same compartment. **i,** CIB1 localization in control and *Tmc5*-KO seminiferous tubules. DAPI, magenta; CIB1, green. **j,** AlphaFold3 model of the TMC5– TMC7–CIB1–CIB4 complex. Scale bars, 10 μm.

To determine whether TMC5 is part of a complex, and identify putative protein partners in vivo, we immunoprecipitated TMC5–mCherry from testis membrane preparations from *Tmc5*–mCherry knock-in mice and evaluated enriched proteins by mass spectrometry (Fig. 5c,d). TMC5–mCherry was the top abundant protein relative to controls, validating the specificity of the affinity purification (Fig. 5c-e and Extended Data Fig. 4a). Notably, TMC7, CIB1, and CIB4 were among the most consistently enriched interactors (Fig. 5c,d). We then validated these interactions biochemically by western blotting TMC5 pull-downs from testis membrane preparations, and confirmed enrichment of both CIB1 and TMC7 in affinity-purified material (Fig. 5e). Reciprocal immunoprecipitation of TMC7-HA from testis membrane preparations similarly recovered TMC5 and CIB1 (Fig. 5f), providing independent biochemical confirmation that TMC5, TMC7, and CIB1 associate within the same molecular complex. Although CIB4 was reproducibly detected in TMC5–mCherry pull-downs by mass spectrometry, available CIB4 antibodies lacked sufficient specificity for reliable western blotting.

To functionally validate the TMC5-CIB1 interaction identified by mass spectrometry, we used a heterologous recruitment assay in COS7 cells^39^. COS7 cells were selected for this assay because they lack endogenous TMC or CIB protein expression, providing a null background in which membrane recruitment can be attributed specifically to the transfected constructs. Additionally, their flat and well-spread morphology makes them well suited to imaging protein distribution between cellular compartments.

We overexpressed both TMC5-mCherry and CIB1-GFP alone and in combination. TMC proteins expressed heterologously are typically aberrantly retained in the endoplasmic reticulum (ER)^40,41^, and we previously showed that TMC5 follows this pattern^17^. When expressed alone, CIB1–GFP was distributed diffusely throughout the cytoplasm (Fig. 5g). By contrast, co-expression with ER-retained TMC5–mCherry caused CIB1-GFP to redistribute to the TMC5-positive compartment (Fig. 5h), supporting an interaction between TMC5 and CIB1. CIB4 followed the same pattern; CIB4-Myc-DDK alone was distributed diffusely through the cytoplasm, but was retained in the ER when co-transfected with TMC5-mCherry (Extended Data Fig. 4b). We could not use this method to confirm TMC5-mCherry and TMC7-Myc-DDK interaction, as TMC7 was also retained in the ER when overexpressed by itself (Extended Data Fig. 4c).

Finally, we examined whether these candidate complex components localize, like TMC5-mCherry, to the spermatid plasma membrane in vivo. We found that CIB1 did indeed localize to the spermatid plasma membrane in control testes (Fig. 5i), but not other cell types which lacked TMC5 expression. Strikingly, this localization was lost in *Tmc5*-KO testes (Fig. 5i), indicating that TMC5 is required for proper CIB1 localization at the spermatid plasma membrane. Together, these data support a model in which TMC5 associates with TMC7, CIB1, and CIB4 in a TMC–CIB complex at the spermatid plasma membrane.

Consistent with this hypothesis, AlphaFold3 models predicted a heteromeric shared molecular assembly containing TMC5, TMC7, CIB1, and CIB4^42^, in which TMC5 and TMC7 form a transmembrane core associated with cytoplasmic CIB proteins (Fig. 5j). In this arrangement, the EF-hand calcium-binding domains of CIB1 and CIB4 are positioned at the cytoplasmic face of the transmembrane core, where they would be ideally placed to sense local Ca^2+^ elevations at the channel mouth, analogous to the regulatory role of CIB proteins in the hair cell mechanotransduction complex^5^.

#### TMC5 loss disrupts CREMτ-dependent transcriptional programs in round spermatids

The cytoskeletal and contractile defects in *Tmc5*-KO spermatids account for many features of the elongation block, but Ca^2+^ signaling also drives gene expression through CREB/CREM-family transcription factors^43^. In the testis, the CREMτ isoform is a master regulator of post-meiotic spermatid differentiation whose transcriptional activity is activated downstream of calcium/cAMP signaling and is essential for spermiogenesis^6^. We therefore asked whether TMC5-dependent Ca^2+^ influx supports expression of CREMτ transcriptional targets in round spermatids, and used single-cell RNA sequencing to assess transcriptome changes across testis cell populations at cell-type-specific resolution.

Consistent with TMC5 being restricted to post-meiotic spermatids, minimal transcriptome changes were observed in spermatogonia and spermatocytes in *Tmc5*-KO testes (Fig. 6a,b and Extended Data Fig. 5a). In stark contrast to controls, MRSs and LRSs from *Tmc5*-KO testes had widely divergent transcriptomes, with later spermatid populations affected more profoundly (Fig. 6a,b and Extended Data Fig. 5b,c).

**Figure 6.**
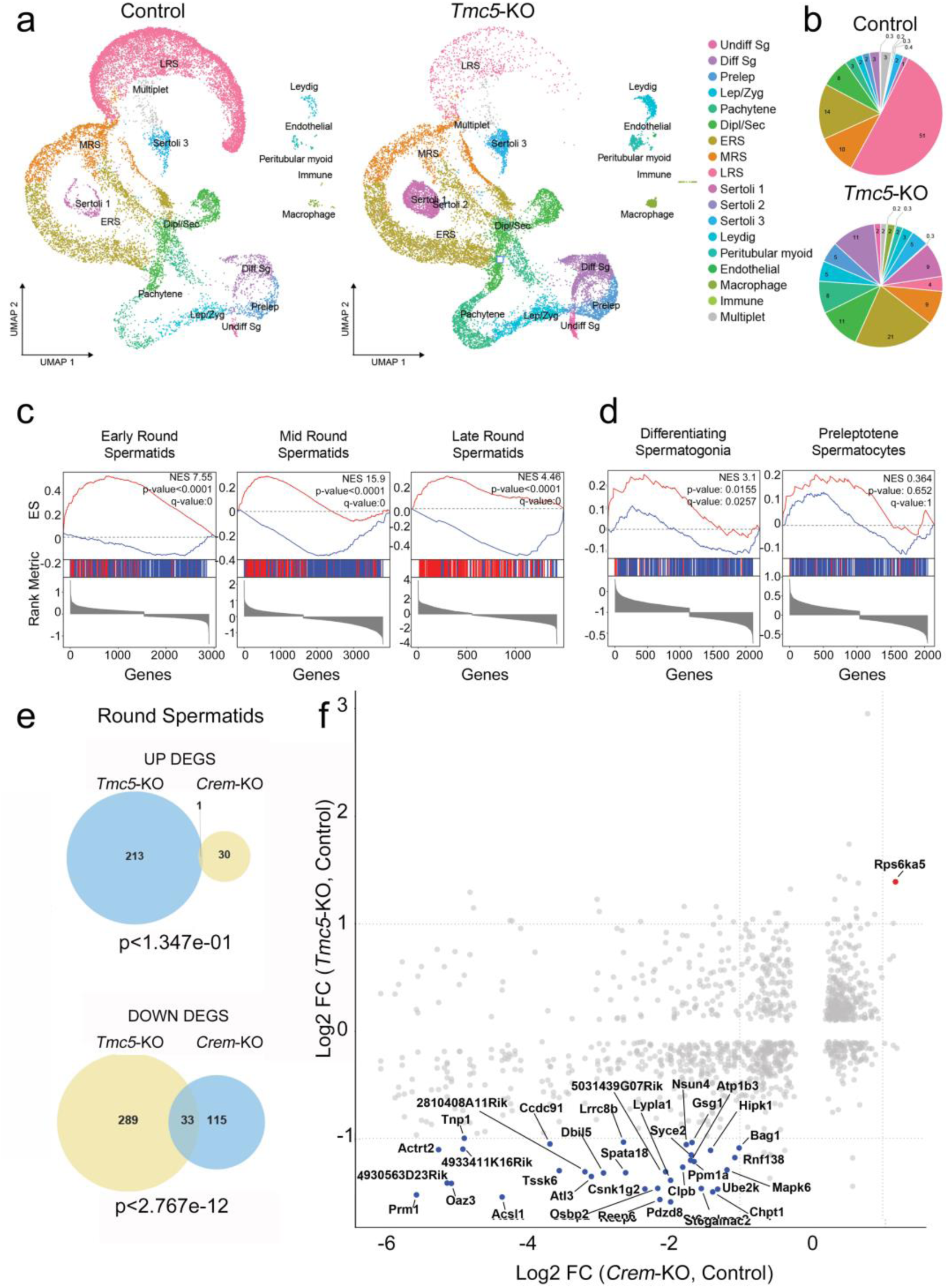
*Tmc5*-KO spermatids exhibit loss of CREMτ-dependent transcription. **a,** UMAP embeddings of single-cell RNA-seq data from control and *Tmc5*-KO testes. Cell populations are color-coded as indicated. **b,** Pie charts showing the proportional representation of each cell type in control and *Tmc5*-KO datasets. **c,** GSEA enrichment plots for direct CREMτ target genes in early, mid, and late round spermatid populations from *Tmc5*-KO relative to control testes. Normalized enrichment scores (NES) and p-values are indicated. **d,** GSEA enrichment plots for CREMτ target genes in differentiating spermatogonia and preleptotene spermatocytes, showing absence of significant enrichment in non-spermatid populations. **e,** Venn diagrams showing overlap between upregulated (top) and downregulated (bottom) differentially expressed genes (DEGs) in round spermatids from *Tmc5*-KO and *Crem*-KO testes. Overlap p-values are shown. **f,** Scatter plot comparing log2 fold changes of round spermatid DEGs in *Tmc5*-KO (y-axis) versus *Crem*-KO (x-axis) relative to control. Blue dots indicate genes concordantly downregulated in both models; selected genes are labeled. Red dots indicate genes concordantly upregulated in both models.

CREMτ is a master transcriptional regulator of spermiogenesis directly activated downstream of calcium/cAMP signaling, but upstream activators of this spermiogenesis gene expression program remain unidentified. Here we asked whether the dysregulated transcripts in *Tmc5*-KO round spermatids overlap with direct CREMτ transcriptional targets. Gene set enrichment analysis revealed strong overlap between genes downregulated in *Tmc5*-KO round spermatids and a curated set of direct CREMτ target genes, defined as genes both bound by CREMτ in published ChIP-seq data^44^ and differentially expressed in *Crem*-KO testes^45^ (Fig. 6c–f and Extended Data Fig. 6a,b). The coordinated reduced expression of these direct CREMτ target genes in *Tmc5*-KO spermatids reveals a requirement for TMC5-dependent Ca^2+^ signaling in activation and/or maintenance of the essential CREMτ-dependent gene expression program during spermiogenesis.

To define whether this relationship was specific to spermatids, we performed a cell-type-resolved comparison of direct CREMτ target genes and genes that were differentially expressed in the *Tmc5*-KO testes. Non-spermatid populations showed little overlap between the two models (Fig. 6d), indicating that TMC5–directed CREMτ transcriptional activation is specific to postmeiotic spermatids.

Altogether, these data position TMC5-dependent mechanosensitive calcium signaling upstream of CREMτ-dependent transcriptional programs in round spermatids, providing a molecular framework linking mechanical stimulation at the spermatid plasma membrane to the stage-specific gene expression required for spermatid elongation and ultimately male fertility.

## Discussion

Here, we show that developing spermatids are mechanoresponsive and convert mechanical stimulation into extracellular Ca^2+^-dependent Ca²⁺ influx. Loss of the predominant mechanically evoked Ca^2+^ response disrupts actomyosin contractility, cytoskeletal organization, and the expression of direct CREMτ transcriptional targets, culminating in failed spermatid elongation and NOA. These findings identify mechanosensitive Ca^2+^ signaling as an essential upstream regulator of critical molecular and cellular events that drive spermiogenesis.

The identification of TMC5 in this process expands the biological scope of the TMC family beyond classical sensory systems. In auditory and vestibular hair cells, TMC1 and TMC2 operate within a highly specialized apparatus, including tip links, LHFPL5, and TMIE^36^, that enables exceptionally rapid and sensitive mechanotransduction. These components are absent from spermatids, where TMC5 instead associates with TMC7, CIB1, and CIB4. The spermatid complex is therefore analogous in organization to sensory TMC–CIB machinery but operates in a distinct molecular and physical context. This distinction suggests that the mechanical properties of TMC-family proteins are substantially shaped by their accessory proteins, membrane environment, and cellular architecture.

Notably, mechanically evoked Ca^2+^ signals increased with spermatid developmental stage and were sustained rather than spike-like. These sustained kinetics of the spermatid Ca^2+^ response may be particularly well suited to the physiology of spermiogenesis. Developing spermatids remain embedded within and mechanically coupled to Sertoli cells while undergoing translocation, nuclear shaping, cytoplasmic remodeling, and large changes in membrane geometry over several days. Continuous remodeling of Sertoli–spermatid adhesions, including the apical ectoplasmic specialization, is likely to expose spermatids to evolving patterns of confinement, adhesion, shear, and membrane tension. A pathway that integrates sustained mechanical loading could coordinate environmental inputs with the gradual remodeling of the spermatid. Although the magnitude, source, and temporal profile of the endogenous forces remain to be defined in intact seminiferous tubules, our findings provide a physiological framework for investigating how those forces regulate germ-cell development

The extent to which TMC5 is itself a directly mechanically gated ion channel remains unresolved. The marked reduction of swelling-evoked Ca^2+^ entry in *Tmc5*-KO spermatids establishes that TMC5 is required for most of the response but does not demonstrate that it forms the conducting pore. TMC5 could instead contribute to the force-sensing apparatus, regulate another channel, or be activated indirectly through accessory proteins or lipid-mediated force transmission. One possibility is that TMC5 responds directly to bilayer tension, resembling OSCA/TMEM63 and TMEM16 proteins, structurally related mechanosensitive proteins that can be activated without a dedicated extracellular force-transmission apparatus and that display relatively high thresholds and slow kinetics^46^. Although TMC5 may share gating properties with OSCA/TMEM63 proteins, *Tmem63a*, *Tmem63b*, and *Tmem63c* transcripts were low or undetectable in spermatids, arguing against these channels as the primary mediators of the response. Directly testing whether TMC5 itself responds to membrane tension will require functional reconstitution of purified TMC5-containing complexes under controlled mechanical stimulation.

The involvement of TMC7 also remains to be defined. While intracellular functions for TMC7 in Golgi homeostasis and acrosome biogenesis have previously been linked to the infertility of *Tmc7*-KO mice^18,19^, TMC7 localization at the spermatid periphery and association with TMC5 suggest an additional role at the plasma membrane. TMC7 may therefore occupy spatially and functionally distinct intracellular and cell-surface pools. Determining whether TMC7 is an obligate component of TMC5-dependent mechanotransduction or performs a separate plasma-membrane function will require compartment-specific functional analysis. Similarly, although CIB1 and CIB4 associate with TMC5, and TMC5 is required for CIB1’s plasma-membrane localization, their precise contributions to force sensing, channel assembly, and Ca^2+^ entry remain to be established.

Canonical mechanically activated channels are unlikely to account for spermatid response to mechanical stimuli. PIEZO channels and TRPV4 were undetectable in postmeiotic germ cells, and spermatids did not respond to PIEZO1 activation. Although TRPV4 functions as a temperature-sensitive channel in mature human sperm^47^, it may be acquired or activated later during epididymal maturation. That *Trpv4*-KO male mice are fertile further argues against an essential role during spermiogenesis^47^. Together, these observations support a mechanotransduction pathway molecularly distinct from those characterized previously in germ cells and sperm.

Consistent with a conserved role in male germ-cell development, TMC5 is expressed in the human testis and has been reported among genes altered in transcriptomic and DNA-methylation studies of teratozoospermia, spermatogenic arrest and low sperm motility^48–50^. Enrichment of TMC5 in both mouse and human spermatids, together with the NOA phenotype of *Tmc5*-KO males, has potential implications for genetically unexplained male infertility. Variants that eliminate reproductive capacity are expected to be rare, highly penetrant, and subject to strong negative selection. They may therefore be poorly represented in population-scale genome-wide association studies but detectable through case-based exome or genome sequencing^20^. TMC5, and potentially TMC7, CIB1, and CIB4, should consequently be considered candidate genes in men with unexplained NOA. Because this machinery is enriched in postmeiotic germ cells, it may also provide a source of selective molecular targets for non-hormonal male contraception.

More broadly, this work identifies plasma-membrane force sensing as an integral component of germ-cell differentiation. It raises the possibility that developing cells in other organs similarly repurpose sensory-like machinery to translate the physical properties of their local environment into cell-type-specific transcriptional and cytoskeletal programs.

## Methods

### Biological samples

Mice from B6SJLF1 and C57BL/6J backgrounds were kept in the animal care facilities at the University of Virginia under controlled temperature and humidity using 12:12-hr light/ dark cycles. Animals were supplied with water and food ad libitum. All animal procedures were approved by the University of Virginia Institutional Animal Care and Use Committee (UVA IACUC: 4382).

### Generation of transgenic mice

Guide RNAs targeting exon 8 of mouse *Tmc5* were designed using the CRISPOR online tool to generate a frameshift-inducing deletion. crRNA, tracrRNA, and Cas9 protein were obtained from Integrated DNA Technologies (IDT). crRNA and tracrRNA were annealed and complexed with Cas9 protein to generate ribonucleoprotein (RNP) complexes before electroporation.

*Tmc5*-KO mice were generated by CRISPR–Cas9-mediated genome editing in C57BL/6J embryos^51^. Female C57BL/6J mice were superovulated and mated with C57BL/6J males, and fertilized single-cell embryos were collected the following day. Embryos were electroporated with the RNP complex using a NEPA21 Super Electroporator (Nepa Gene Co.). Electroporated embryos were cultured overnight in KSOM medium at 37°C in 5% CO₂ and transferred at the two-cell stage into the oviducts of pseudopregnant ICR females (Inotiv). Electroporation and embryo transfer procedures were performed by the University of Virginia Genome Engineering and Modeling (GEMM) Core. Founder animals were identified by PCR genotyping and Sanger sequencing of tail-derived DNA, which confirmed a 44-bp deletion in exon 8 of *Tmc5* predicted to produce a frameshift and loss-of-function allele. The founder mouse was backcrossed to C57BL/6J mice for 6 generations, to minimize the likelihood of off-target mutations in the experimental mice. Heterozygous (*Tmc5*^+/–^) littermates were used as controls throughout this study.

### Spermatogenic cells isolation from mouse testis

Collagenase A (50 mg/mL) and DNase I (20 mg/mL) stocks were prepared in HBSS, stored at 4°C before use and at −80°C thereafter, and aliquoted to minimize freeze–thaw cycles. Digestion media per mouse were freshly prepared as follows: digestion medium I (F12 supplemented with collagenase A (0.1 mL) and DNase I (0.1 mL), total 10 mL); 5% Percoll solution (HBSS (10×, 4 mL), Percoll (2 mL), and ddH₂O (34 mL), total 40 mL); and digestion medium II (F12 supplemented with trypsin (0.25%, 2 mL) and DNase I (0.1 mL), total 10 mL).

Testis single-cell suspensions were prepared as previously described with modifications^52^. Briefly, testes were collected from sacrificed mice, and the tunica albuginea were removed. Decapsulated testes were incubated in digestion medium I (10 mL) at 37°C for 5 min, mechanically dissociated by gentle pipetting, and further incubated for 5 min with intermittent pipetting until seminiferous tubules were released while remaining structurally intact. The digested tissue was layered onto 5% Percoll (40 mL) at room temperature (20-25°C) and allowed to separate for 5 min, during which interstitial cells remained in the supernatant, and seminiferous tubules sedimented. The upper 35 mL was removed, and the seminiferous tubules were collected and transferred to digestion medium II (10 mL).

Secondary digestion was performed at 37°C for 15 min with gentle pipetting until tubules were no longer visible. Trypsin was neutralized with 3 mL fetal bovine serum. The suspension was sequentially filtered through 70- and 40-μm strainers and centrifuged at 500 × g for 10 min at 4°C.

### Spermatogenic synchronization

Spermatogenesis was synchronized using WIN 18,446 and RA, adapted from the protocol previously described^53^ with modifications to improve survival of neonatal inbred mice. Neonatal male mice were orally administered WIN 18,446 (Cayman Chemical, Item No. 14018) once daily from P2 through P10. WIN 18,446 was administered at 60 μg/g body weight, corresponding to 60% of the 100 μg/g body-weight dose used in the published protocol. This reduced dose and modified treatment schedule were used to decrease treatment-associated mortality while maintaining suppression of endogenous RA-dependent spermatogonial differentiation. On P11, mice received a single subcutaneous injection of 100 μg RA (Sigma-Aldrich, R2625) dissolved in 10 μL DMSO to release the WIN 18,446-induced differentiation block and synchronously initiate spermatogonial differentiation. Following RA administration, no additional WIN 18,446 treatment was given, and mice were allowed to progress through spermatogenesis. For stage-specific experiments, LRS were identified and used for experiments at P32–P33, whereas ERS were identified and used for experiments at P37 when both second wave ERS and first wave ES are present.

### Fura-2 imaging and hypotonic mechanical stimulation

Fura-2 AM stock solution (500 μM) was prepared by dissolving 50 μg Fura-2 AM (Invitrogen; MW 1001.86 g/mol) in 99.8 μL DMSO and stored in aliquots to minimize freeze–thaw cycles. Isotonic and hypotonic buffers were prepared as previously described. The isotonic buffer (∼314 mOsm, pH 7.4) contained NaCl (96 mM), KCl (5.3 mM), MgSO₄ (0.8 mM), NaH₂PO_4_ (1 mM), HEPES (5 mM), and mannitol (100 mM), with either Ca-acetate (1.8 mM) or EGTA (1.8 mM) to generate calcium-containing or calcium-free conditions, respectively. The hypotonic buffer (∼214 mOsm, pH 7.4) was identical in composition except that mannitol was omitted to reduce osmolarity. All buffers were adjusted to pH 7.4, and osmolarity was confirmed before use. Cells were resuspended at a density of 6 × 10^6^ cells/mL and loaded with 5 μM Fura-2 AM in suspension under isotonic conditions at room temperature for 45 min. Following loading, cells were washed by centrifugation to remove excess dye.

For imaging, 384-well non-binding, flat-bottom μclear microplates (Greiner Bio-One, Cat. No. 781906) were coated with Cell-Tak (Corning) according to the manufacturer’s instructions. A total of 10 μL of the cell suspension (∼6 × 10^4^ cells per well) in isotonic buffer, with or without calcium as indicated, was added to each well. Plates were sealed with tape to prevent evaporation and spun briefly to allow cell attachment. Ratiometric calcium imaging was performed by sequential excitation at 340 nm and 380 nm, and intracellular calcium levels were quantified as the fluorescence ratio (F_340_/F_380_). Fura-2 emission was collected at 510 nm using a DG4 illuminator (Sutter Instruments) for excitation and an ORCA-Flash 4.0 V2 CMOS camera (Hamamatsu) controlled by SlideBook 6 software for image acquisition. Cells were imaged at 10-s intervals throughout the experiment. Hypotonic stimulation was induced during imaging by the addition of a 90 μL hypotonic buffer directly to each well, and changes in intracellular calcium were monitored in real time. Ionomycin was added at the end of each recording as a positive control for calcium influx. For trace analysis, responses were expressed as Δ(F_340_/F_380_). Baseline F_340_/F_380_ ratio was defined as the mean ratio over the first 20 frames (100s) of each recording. Δ(F_340_/F_380_) was calculated by subtracting this baseline value from the F_340_/F_380_ ratio at each subsequent time point. Peak Δ F_340_/F_380_ (Fig. 1g) was defined as the maximum value reached during the stimulation window. Following imaging, AO/PI staining (Revvity) was performed for cell identification and live/dead cell assessment.

### Chemical stimulation of cells

For chemical activation of PIEZO1, cells were stimulated with 50 μM Yoda1 (Cayman Chemical, Item No. 21904). Cells were stimulated with the calcium ionophore A23187 (calcimycin; Abcam, ab120287) to induce an increase in intracellular Ca^2+^. A23187 was prepared as a 10 mM stock solution in DMSO and diluted into the experimental solution to a final concentration of 10 μM immediately before use.

### Single cell library preparation and single cell RNA sequencing

The generation of single cell emulsions was performed by the School of Medicine Genome Analysis and Technology Core, RRID:SCR_018883, using the 10x Genomics Chromium X platform and the Chromium GEM-X Single Cell 3’ Chip Kit (P/N 1000690, 1000691, 1000215). Barcoded cDNA was generated from biological replicates (KO and control) using the manufacturer protocol (10x Genomics) and assessed for quality using the Agilent 4200 TapeStation Instrument and the Agilent D5000 HS kit. Indexed single cell libraries were prepared from the cDNAs and sizing performed on the TapeStation as above. Fluorometry (Qubit 3) was used to determine the concentration of each sample which were subsequently pooled using an equimolar ratio for a QC run on the Illumina NextSeq2000 using a P1-100 cycle kit targeting ∼25,000 reads/cell. FastQ files were generated using Illumina’s BCLConvert for downstream data analysis. FASTQ files were aligned to the mm10-3.0.0 mouse reference genome, and barcode filtering and UMI counting were performed using Cell Ranger v9.0.1 (10X Genomics) *mkfastq* and *count* pipelines with default parameters. Raw and processed data were deposited in GEO under accession number GSE342049.

### Single-cell RNA sequencing analysis

All analyses were performed in R (v4.4.1)^54^ using Seurat (v5.5.1)^55^. Ambient RNA contamination was corrected using SoupX (v1.6.2)^56^, and dataset integration was performed using Harmony (v2.0.5)^57^. Additional analyses were performed using Monocle3 (v1.4.26)^58–60^, SeuratWrappers (v0.4.0)^61^, and the tidyverse suite of packages (v2.0.0)^62^. Visualization was performed using ggplot2 (v4.0.3)^63^, and EnhancedVolcano (v1.24.0)^64^.

### Quality Control and Filtering

For each sample, ambient RNA-corrected count matrices were imported into Seurat using *CreateSeuratObject()* with minimum thresholds of three cells per gene and 200 detected genes per cell. The percentage of mitochondrial transcripts was calculated using *PercentageFeatureSet()*. Quality control (QC) assessment was performed independently for each sample using an interquartile range (IQR)-based approach. Lower and upper thresholds for gene counts (nFeature_RNA) and transcript counts (nCount_RNA) were determined as Q1 - 1.5 times IQR and Q3 + 1.5 times IQR, respectively, with a minimum detection of 200 genes per cell. Cells with more than 10% mitochondrial transcripts were excluded.

### Data Integration and Dimensionality Reduction

Filtered Seurat objects were merged into a single dataset using Seurat’s merge() function. Data were normalized using *NormalizeData()*, highly variable genes were identified using *FindVariableFeatures()*, scaled using *ScaleData(*), and principal component analysis was performed using RunPCA(). Batch effects between samples were corrected using Harmony integration in PCA space. Cell-to-cell neighborhoods were identified using *FindNeighbors()* on the first 30 Harmony dimensions, followed by graph-based clustering using *FindClusters()* (res = 1). Two-dimensional visualization was performed using Uniform Manifold Approximation and Projection (UMAP). Cell populations were annotated based on expression patterns of established testicular germ cell and somatic cell markers, as described^13^. Marker expression was evaluated using UMAP feature plots *(FeaturePlot)* and violin plots *(VlnPlot)* across clusters and experimental conditions.

### Differential Gene Expression Analysis

Following dataset integration and cluster annotation, layers were rejoined using *JoinLayers()* to restore unified count and expression matrices. Cells were grouped according to annotated *cell* populations, including undifferentiated spermatogonia (Undiff Sg), differentiating spermatogonia (Diff Sg), preleptotene spermatocytes (Prelep), leptotene/zygotene spermatocytes (Lep/Zyg), pachytene spermatocytes, diplotene/secondary spermatocytes (Dipl/Sec), early round spermatids (ERS), mid-round spermatids (MRS), late round spermatids (LRS), Sertoli cells, Leydig cells, peritubular myoid cells, endothelial cells, macrophages, and immune cells. For each cell type, cells were subset from the integrated Seurat object and reassigned according to genotype (Control or KO). Differential expression between genotypes was performed independently within each cell population using Seurat’s *FindMarkers()* function with the Wilcoxon rank-sum test and a minimum expression threshold of 25% of cells.

To compare transcriptional changes between *Tmc5*-KO and a previously published CREM knockout dataset^45^, differential expression results were matched based on shared genes. Scatter plots were generated using ggplot2, each point representing a gene detected in both datasets. The x-axis represents the log_2_ fold-change from the CREM knockout dataset, while the y-axis represents the log_2_ fold-change from the *Tmc5*-KO dataset. Comparisons were performed for early round spermatids (ERS), mid-round spermatids (MRS), late round spermatids (LRS), and a combined round spermatid (RS) dataset generated by pooling ERS, MRS, and LRS genes. Reference lines show log2 fold-change values of ±1. Based on their log_2_FC values across both datasets, genes were classified into three response types: shared upregulation (>1), shared downregulation (<-1), or oppositely regulated.

To determine whether overlap between TMC5 and CREM-regulated gene sets exceeded random expectation, hypergeometric enrichment analyses were performed. Significant genes were defined as those with *P < 0.05*, while upregulated and downregulated genes were defined by log_2_ fold-change thresholds greater than 1 or less than -1, respectively. Hypergeometric tests were conducted for all DEGs, upregulated DEGs, and downregulated DEGs within ERS, MRS, LRS, and pooled RS datasets. For each comparison, the gene universe was defined as the intersection of genes detected in the corresponding CREM and TMC5 datasets. The probability of observing the measured overlap by chance was calculated using the cumulative hypergeometric distribution *(phyper)*. Gene overlaps were visualized using Venn diagrams generated with the *VennDiagram* package.

Differential Gene Set Enrichment Analysis (DGSEA) was also performed to compare transcriptional signatures between *Tmc5*-KO and the CREM-target dataset^45^. Genes significantly upregulated (log2FC > 0, *P < 0.05*) or downregulated (log2FC < 0, *P < 0.05*) in CREM-targets were used as reference gene sets. For each TMC5 cell population, genes were ranked by log2 fold change and analyzed using the *dgsea_targeted()* function with weighted enrichment statistics and 10,000 permutations. Hallmark gene sets from MSigDB (*Mus musculus*) were included as background pathways. Enrichment results were visualized using DGSEA mountain plots generated with make_mountain_plots().

### Pseudotime Trajectory Analysis

For the pseudotime analysis, germ-cell populations were extracted from the integrated Seurat dataset. Control and *Tmc5*-KO datasets were analyzed separately. Seurat objects were converted to Monocle3 cell data sets using *as.cell_data_set()*. Cell-type annotations were assigned as cluster identities, and all cells were assigned to a single partition to enable construction of a continuous developmental trajectory. Principal graphs were learned using *learn_graph()* with partitioning disabled. Cells were ordered along pseudotime using *order_cells()*, with undifferentiated spermatogonia specified as the root population. Pseudotime trajectories were visualized using *plot_cells()*, and Control and *Tmc5*-KO trajectories were displayed using a common pseudotime color scale to facilitate comparison of developmental progression.

#### Antibodies and other reagents

Primary antibodies used in this study included anti-TMC7 (Sigma, HPA029465), anti-CIB1 (Sigma, HPA042413), anti-acetylated α-tubulin (Sigma, T6793), anti-HA (Cell Signaling, #3724), anti-mCherry (Invitrogen, M11217), and anti-pMLC2 (Cell Signaling, #3671). For immunofluorescence staining, Alexa Fluor-conjugated secondary antibodies included anti-rabbit Alexa Fluor 488 (Invitrogen, A48282), anti-rabbit Alexa Fluor 568 (Invitrogen, A21069), and anti-mouse Alexa Fluor 568 (Invitrogen, A11019). For western blot analyses, IRDye-conjugated secondary antibodies included anti-rat 680 (Jackson ImmunoResearch, 112-625-167), anti-rabbit 790 (Jackson ImmunoResearch, 111-655-144), and anti-mouse 680 (Jackson ImmunoResearch, 115-625-146). Additional reagents included peanut agglutinin (PNA)-lectin conjugate (Invitrogen, L21409) and Hoechst dye (Invitrogen, H3570).

#### Membrane isolation

Testes were mechanically homogenized in an ice-cold dilution buffer containing 50 mM Tris-Cl (pH 7.6), 50 mM NaCl, and 5 mM EDTA, supplemented with cOmplete™ Protease Inhibitor Cocktail (Roche, Cat. No. 11836170001) using a Waring commercial blender. Following homogenization, samples were subjected to low-speed centrifugation at approximately 12,500 × g for 20 min at 4°C to remove cellular debris and unbroken material. The resulting supernatant was collected and further centrifuged at approximately 200,000 × g for 2 hr at 4°C to isolate membrane fractions^65^.

### Co-immunoprecipitation and western blot analysis

Testes were collected and flash-frozen prior to use. Whole testis or membrane fractions were used for the pulldown. Frozen tissue was mechanically homogenized and lysed at 4°C for 2 hr in lysis buffer, dilution buffer with 2% (w/v) n-Dodecyl-β-D-maltoside (DDM). Lysates were clarified by ultracentrifugation at 186,000 × g for 50 min at 4 °C, and the supernatant was collected and diluted with a detergent-free buffer to reduce the DDM concentration.

Magnetic beads tagged with mCh nanobody (proteintech, ChromoTek RFP-Trap® Magnetic Agarose) were equilibrated by washing three times in a dilution buffer lacking DDM. Clarified lysates were then incubated with equilibrated beads (50 μL bead slurry per 1 mL lysate) for 2 hr at 4°C with end-over-end rotation. Beads were collected magnetically and washed three times with a wash buffer containing 20 mM Tris-Cl (pH 7.6), 150 mM NaCl, and cOmplete™ Protease Inhibitor Cocktail (Roche) to remove non-specifically bound proteins.

Bound proteins were eluted in 2X Laemmli SDS sample buffer (Bio-Rad) by heating at 50°C for 10 min. Eluted proteins were separated by SDS–PAGE and transferred onto membranes for western blot analysis. Membranes were blocked in 5% skim milk for 1 hr at room temperature or overnight at 4°C, followed by incubation with primary antibodies diluted in Solution 1 (Can Get Signal) overnight at 4°C or for 2 hr at room temperature. After washing, membranes were incubated with HRP-conjugated secondary antibodies for 45 min at room temperature. Protein signals were detected using chemiluminescence

### Mass spectrometry

Just prior to digestion, the samples containing magnetic beads were thawed and 150 μL of 50 mM ammonium bicarbonate was added. Samples were gently agitated to mix. Next, 1 μL of dithiothreitol (DTT) was added to each tube and 1 μg of trypsin (MS-grade Trypsin Gold, Promega, V5280) was added for proteolytic digestion. The samples were incubated at 37°C with gently shaking for 16 hr. Samples were then magnetized and supernatant containing peptides was removed and saved to new low-bind Eppendorf tubes. Beads were washed with 75 μL water, magnetized, and supernatant added to the collection tubes. Finally, 1 μL of concentrated formic acid was added to each collection tube to stop trypsin activity. Samples were desalted using Pierce desalting spin columns (Thermo Fisher Scientific, 89852), the eluent dried via Speedvac and reconstituted with 0.1% formic acid to 1 μg/μL final concentration. Desalted samples were analyzed in triplicate by nanoLC-MS/MS using a Dionex Ultimate 3000 (Thermo Fisher Scientific, Bremen, Germany) coupled to an Orbitrap Eclipse Tribrid mass spectrometer (Thermo Fisher Scientific, Bremen, Germany).

Each injection of 1 μg was loaded onto an Acclaim PepMap 100 trap column (300 μm × 5 mm × 5 μm C18) and gradient-eluted from an Aurora Ultimate TS analytical column (75 μm × 25 cm, 1.7 μm C18) equilibrated in 100% solvent A (0.1% formic acid in water) and 0% solvent B (80% acetonitrile in 0.1% formic acid). The peptides were eluted into the mass spectrometer at 400 nL/min up to 100% B over a period of 97 min.

MS2 spectra were acquired using a data-dependent instrument method with the following settings: positive ion mode was used with 1.6 kV at the spray source, RF lens at 30% and data dependent MS/MS acquisition with XCalibur version 4.3.73.11. Full MS scans were acquired in the Orbitrap from 375 to 1500 m/z with 120,000 resolution. Data dependent selection of precursor ions was performed in Cycle Time mode, with three seconds in between Master Scans, using an intensity threshold of 2 × 10^4^ ion counts and applying dynamic exclusion (n = 1 scans within 30 s for an exclusion duration of 60 s and ±10 ppm mass tolerance). Monoisotopic peak determination was applied, and charge states 2–8 were included for HCD MS2 scans (quadrupole isolation mode; 1.6 m/z isolation window, normalized collision energy at 30%). The resulting fragments were detected in the Orbitrap at 15,000 resolution with normalized AGC target of 100% and dynamic maximum injection time mode.

Raw MS data was searched using Proteome Discoverer against a custom fasta database that contained all mouse Gencode vM35 protein sequences as well as mCherry sequences. The following parameters were used: trypsin with maximum 2 missed cleavage sites, 10 ppm precursor mass tolerance and 0.02 Da fragment ion mass tolerance, dynamic amino acid modifications (Oxidation / +15.995 Da (M), Phospho / +79.966 Da (S, T, Y), Carbamidomethyl/ +57.021(C)) and dynamic n-terminal modifications (Acetyl / +42.011 Da, Met-loss / -131.040 Da (M), and Met-loss+Acetyl / - 89.030 Da (M))^66^.

### Immunofluorescence staining and imaging

Mouse testes and epididymides were fixed in 4% paraformaldehyde (PFA) at 4°C for 24– 48 hr, cryoprotected in sequential 10%, 20%, and 30% sucrose solutions, embedded in OCT compound, and stored at -80°C. Frozen testes were cryosectioned at 20 μm thickness using a cryostat maintained at -20°C.

For immunofluorescence staining, tissue sections were washed in PBS, permeabilized in 0.5% PBST, and subjected to antigen retrieval using 10 mM sodium citrate buffer (pH 6.0) at 95-100°C for 20 min. Sections were then blocked in 10% normal goat serum (NGS, Invitrogen) for 2 hr at room temperature or overnight at 4°C and incubated with primary antibodies overnight at 4°C, followed by Alexa Fluor-conjugated secondary antibodies for 45 min at room temperature in the dark. Alexa Fluor-conjugated phalloidin staining was performed where indicated. Slides were mounted using Fluoromount-G™ Mounting Medium with DAPI (Invitrogen).

Confocal imaging was performed using a CSU-W1 spinning disk (Yokogawa) confocal microscope system. Images were acquired at a sampling frequency of 30-60 nm at the camera sensor using NIS-Elements software (Nikon Instruments)^67^.

#### Histology and seminiferous tubule staging

Testes were fixed in either Bouin’s solution or 4% PFA overnight at 4°C and then washed several times in cold 1× PBS. Afterwards testes fixed in Bouin’s solution were processed for embedding in paraffin using standard methods while those fixed in PFA were prepared for embedding in O.C.T. and subsequent cryosectioning as previously described^68^. Bouin’s-fixed and paraffin-embedded 5 µm sections were stained with period acid Schiff using standard methods and stages of the seminiferous epithelium determined based on specific germ cell associations^69^ and aided by visualizing the developing acrosome.

### Statistics

Statistical analyses were performed using GraphPad Prism (GraphPad Software). Data are presented as mean ± SEM unless otherwise indicated. Comparisons between two independent groups were performed using two-tailed *t*-tests. Comparisons among three or more groups were performed using ordinary one-way ANOVA followed by multiple-comparisons test. Mixed-effects analysis was used for repeated or matched datasets where appropriate. A *P* value < 0.05 was considered statistically significant. The statistical test used for each experiment, sample size (*n*), and corresponding *P* values are indicated in the figure legends.

## Data availability

The single-cell RNA sequencing data generated in this study have been deposited in the Gene Expression Omnibus (GEO) under accession number GSE342049 (https://www.ncbi.nlm.nih.gov/geo/query/acc.cgi?acc=GSE342049). To review this dataset prior to publication, navigate to the link above and enter reviewer token mvofsqwybrmzjal into the box. The mass spectrometry proteomics data have been deposited to the ProteomeXchange Consortium via the PRIDE partner repository under dataset identifier PXD081547.

## Acknowledgements

We thank Daniel Grigsby and the UVA Genetically Engineered Murine Model Core (RRID:SCR_025473) for generating The *Tmc5*-KO and *Tmc7*-HA mice used in this study. We thank Katia Sol-Church and Alyson Prorock of the UVA Cancer Center Genomics Core (RRID:SCR_018883) for Single cell library preparation and Single cell RNA-seq. We thank Julia Kachar for assisting with biochemical isolation of TMC5-complex from spermatid membranes. The authors thank Armando Rodriguez of the UT San Antonio Research Computing Support Group for computational support. This study utilized the ARC high-performance computing cluster operated by UT San Antonio Enterprise Research Solutions.

## Author Contributions

S.E. conceived and supervised the study and wrote the manuscript. W.S.Z. performed and analyzed the majority of the experimental work in the study, including all experiments not otherwise attributed below, and contributed to experimental design, data interpretation, figure preparation, and manuscript editing. J.D. established Fura-2 ratiometric calcium imaging in the Desai laboratory and trained W.S.Z. in its use. B.N.D. provided the calcium-imaging facility and expertise in ratiometric calcium imaging and ion-channel physiology. C.M.S. and M.K. provided technical and conceptual guidance on hypotonic-stimulation calcium-imaging experiments and their interpretation. J.S. generated the Tmc5-KO mice, including CRISPR target design, founder screening, and backcrossing onto the C57BL/6J background. E.J. and G.S. performed proteomic analyses and interpreted the proteogenomic data, and E.J. helped edit the manuscript. L.R.C. and B.P.H. analyzed the single-cell RNA-sequencing results, and S.S.B. contributed to the analysis and interpretation of transcriptomic data. B.C. performed western blotting of human and mouse tissues and histological analyses of wild-type and Tmc5-KO testes, and C.B.G. contributed to the interpretation of these experiments, conceptual discussions, and editing of the manuscript. L.S. and F.L. performed protein pull-down experiments and spermatogenic synchronization experiments. A.W. contributed to revision of the manuscript. All authors reviewed and approved the final manuscript.

---

**Extended Data Fig. 1.**
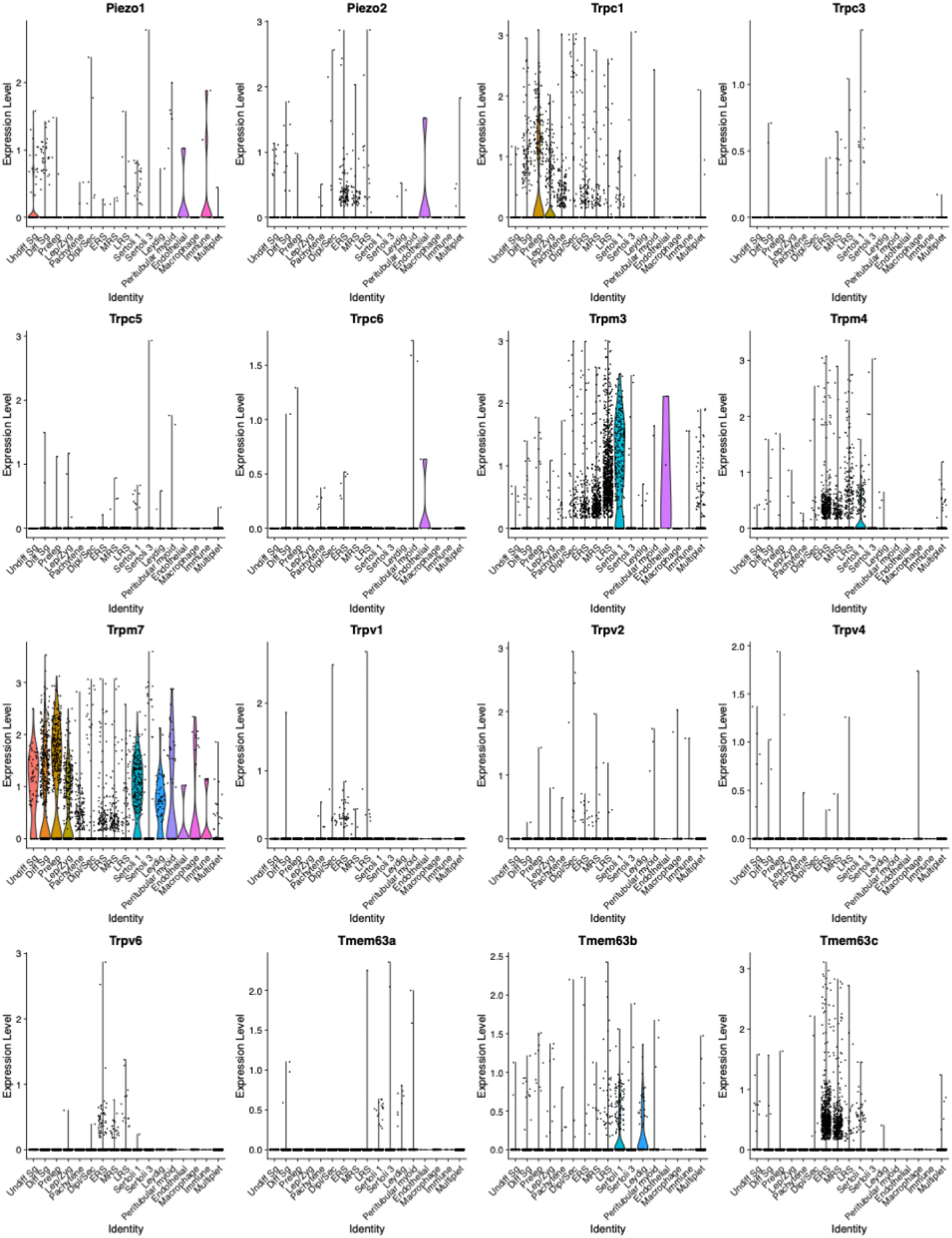
Single-cell RNA-sequencing analysis of canonical mechanosensitive channel expression in testes. Violin plots show the distribution of normalized transcript expression across annotated testicular germ-cell and somatic-cell populations for Piezol, Piezo2, Trpcl, Trpc3, Trpc5, Trpc6, Trpm3, Trpm4, Trpm7, Trpvl, Trpv2, Trpv4, Trpv6, Tmem63a, TmemeSb, and Tmem63c.

**Extended Data Fig. 2.**
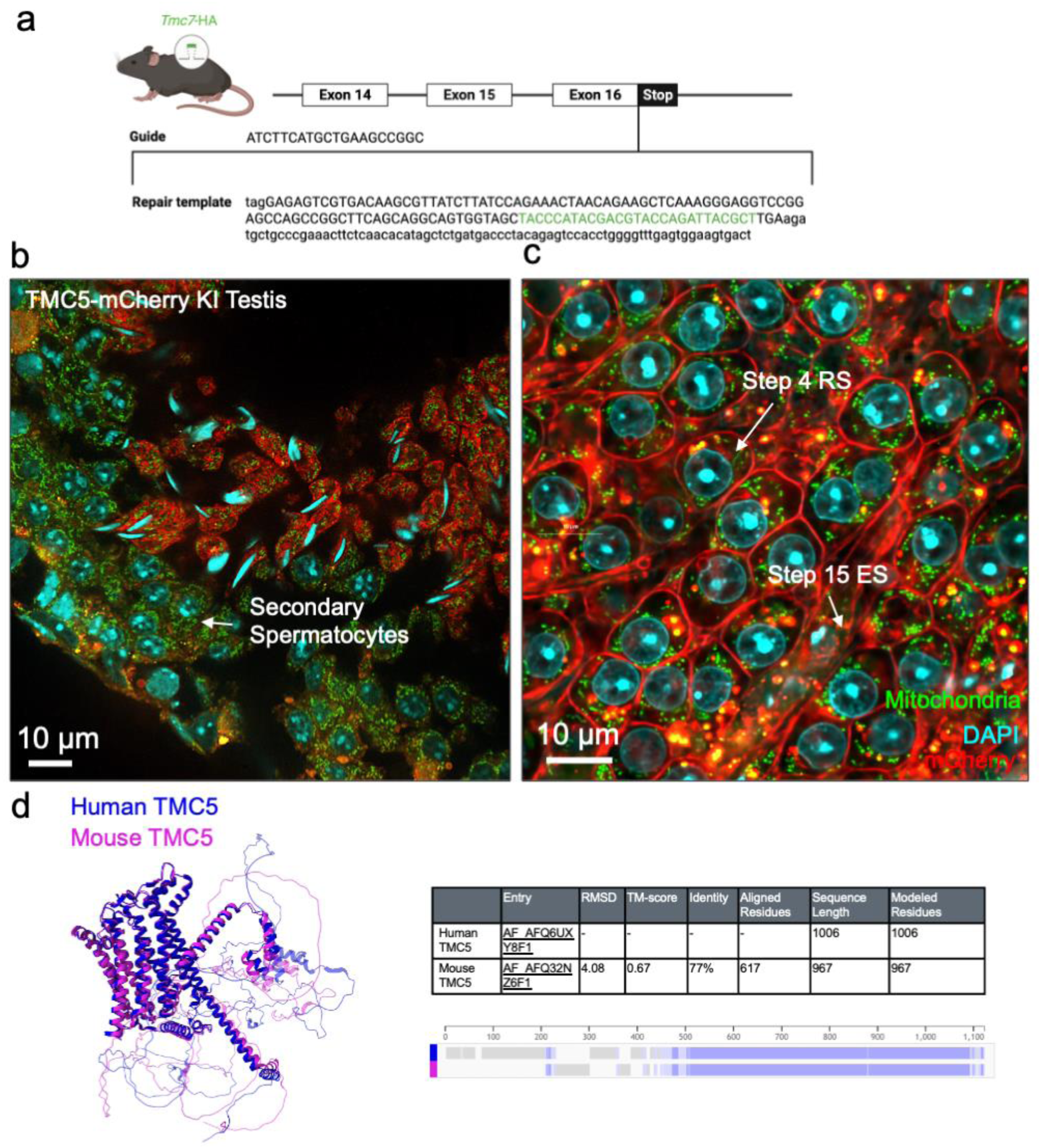
Conservation and developmental expression of TMC5 and TMC7 in mouse spermatogenesis, **a**, Schematic of the CRISPR/Cas9 strategy used to generate the Tmc7-HA knock-in mouse line. The guide RNA targets the region near the stop codon in exon 16, and the repair template introduces an in-frame HA epitope sequence at the C-terminus of TMC7. **b,** Representative fluorescence image of a 7mc5-mCherry *knock-in* testis section showing no detectable TMC5-mCherry signal in secondary spermatocytes, **c,** Higher-magnification image showing the onset of TMC5-mCherry expression in live step 4 round spermatids and its persistence in step 15 elongating spermatids. TMC5-mCherry is shown in red, mitochondria in green, and DAPI-stained nuclei in cyan, d, Structural superposition of predicted human TMC5 (blue) and mouse TMC5 (magenta), together with sequence and structural alignment metrics. Human and mouse TMC5 share 77% sequence identity across 617 aligned residues.

**Extended Data Fig. 3.**
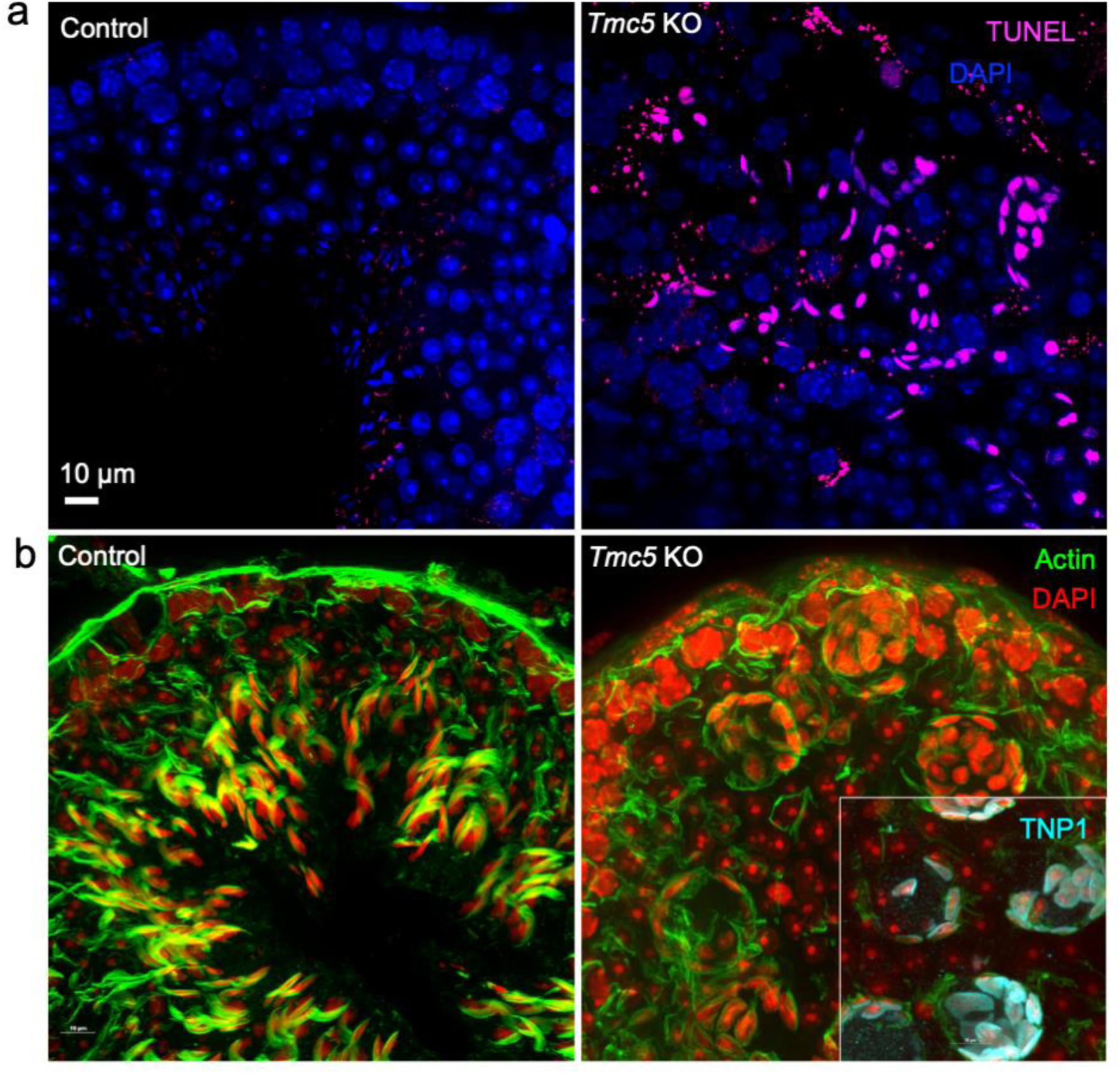
Increased apoptosis in *Tmc5-K0* testes in elongating spermatids, **a,** Representative seminiferous tubule sections from control and *Tmc5-K0* testes stained by TUNEL to detect apoptotic cells. TUNEL-positive cells are shown in magenta and nuclei are counterstained with DAPI (blue). Increased TUNEL labeling is observed in *Tmc5-K0* tubules, **b,** Representative control and Tmc5-K0 seminiferous tubules stained for F-actin (green) and DAPI (red), showing no overt morphological defects in round spermatids but in elongating spermatids. The inset shows TNP1-positive elongating spermatids (cyan) within Tmc5-K0 tubules.

**Extended Data Fig. 4.**
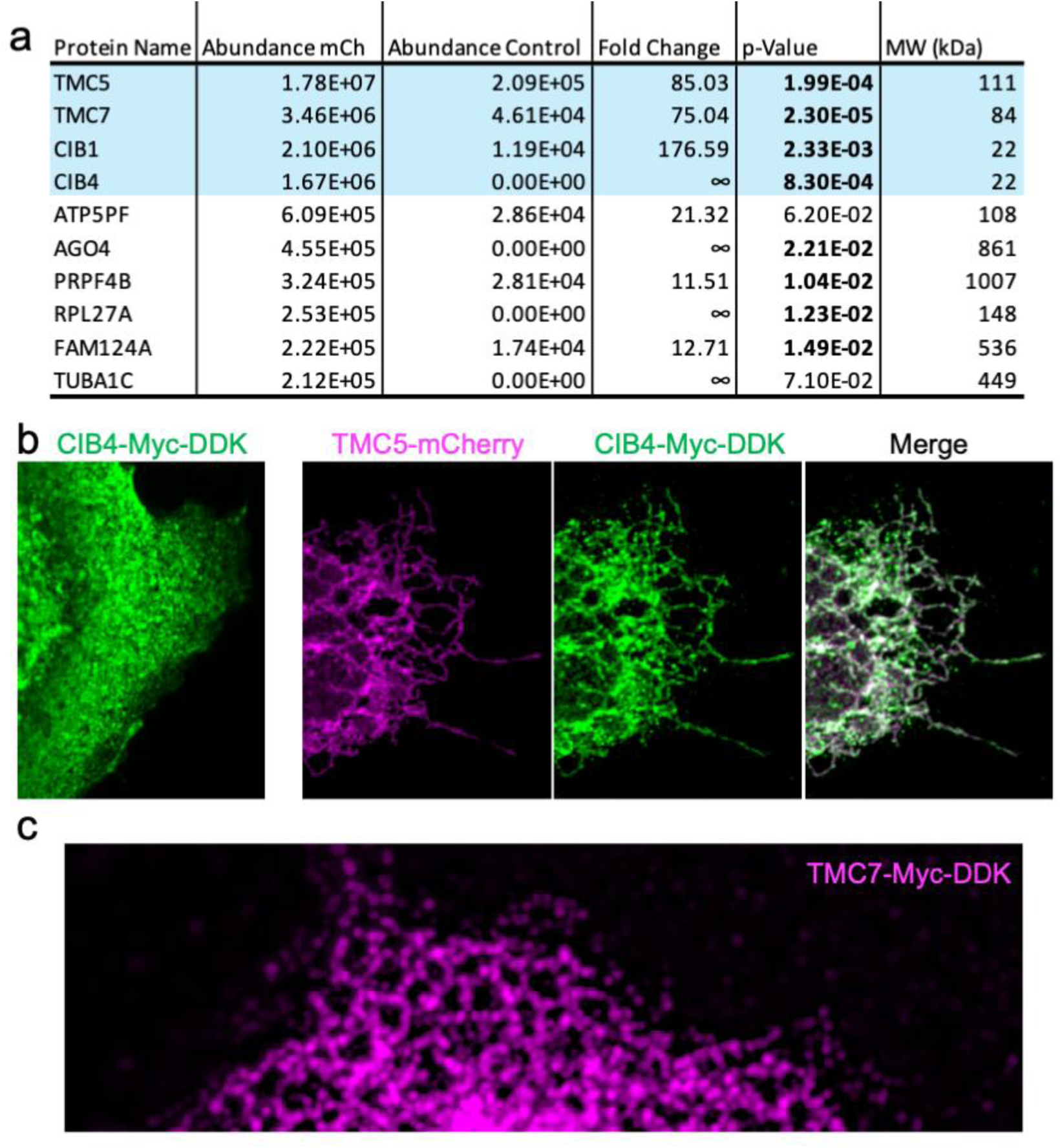
Identification and validation of candidate TMC5-interacting proteins, **a,** Proteins enriched by TMC5-mCherry affinity purification followed by mass spectrometry. Protein abundance in the TMC5-mCherry and control samples, fold enrichment, statistical significance, and predicted molecular weight are shown. TMC5, TMC7, CIB1, and CIB4 were among the most highly enriched candidates, **b,** Representative fluorescence images of COS7 cells expressing CIB4-Myc-DDK alone, and co-expressing TMC5-mCherry (magenta) and CIB4-Myc-DDK (green). The merged image shows substantial colocalization of TMC5 and CIB4 in the ER. **c,** Representative fluorescence image showing the ER localization of TMC7-Myc-DDK in transfected COS7 cells.

**Extended Data Fig. 5.**
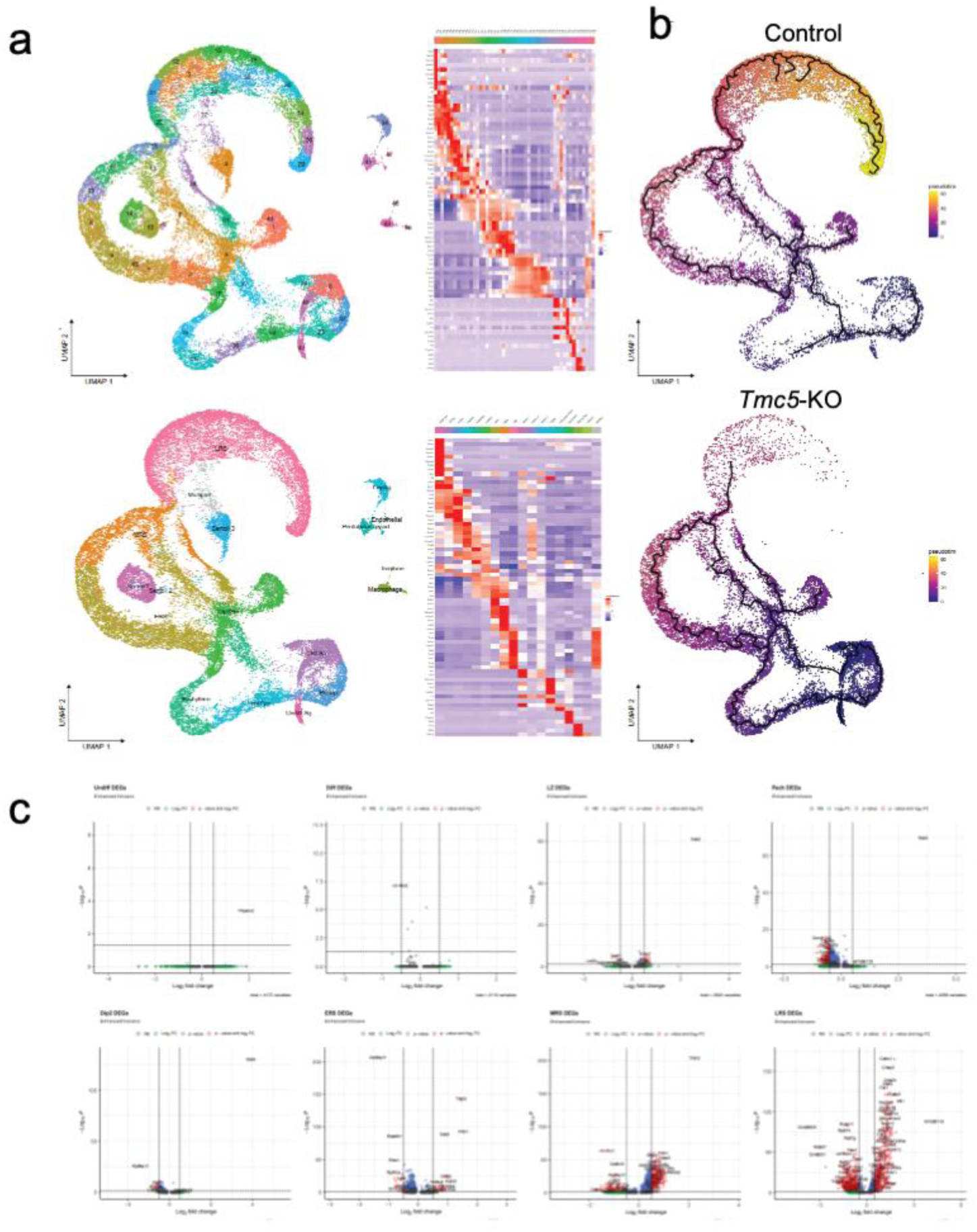
Single-cell RNA-sequencing analysis of Control and *Tmc5* KO testes. **a**, UMAP projections of all cells from Control and KO testes, grouped according to similarities in gene-expression profiles. Heatmaps show the expression of representative marker genes used to identify and annotate cell populations, **b,** Inferred pseudotime trajectories projected onto the Control and 7mc5-KO UMAP embeddings. Colors indicate relative pseudotime, and black lines represent the inferred developmental paths through spermatogenic differentiation, c, Volcano plots showing differential gene expression between Control and 7mc5-KO cells within undifferentiated spermatogonia, differentiating spermatogonia, leptotene/zygotene spermatocytes, pachytene spermatocytes, diplotene spermatocytes, and early, mid, and late round spermatids. Dashed lines indicate the statistical-significance and fold-change thresholds.

**Extended Data Fig. 6.**
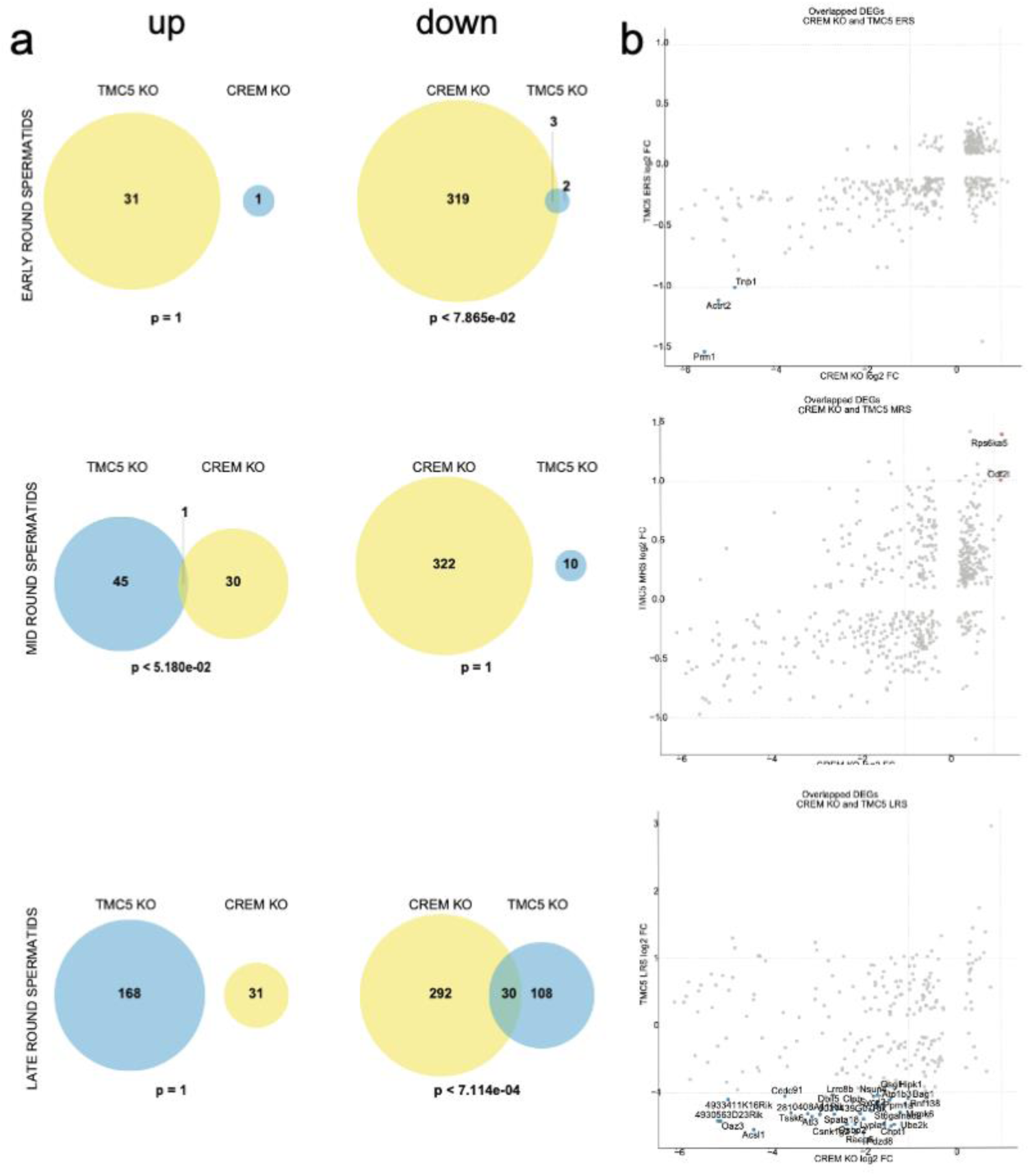
Comparison of differentially expressed genes in *Tmc5-* and *Crem-KO* round spermatids, **a,** Venn diagrams showing the overlap between differentially expressed genes identified in 7”mc5-KO and Crem-KO early, mid, and late round spermatids, **b,** Scatter plots comparing log_2_ fold changes for genes differentially expressed in both ***Tmc5-*** KO and Crem-KO early, mid, and late round spermatids.

## References

1. O’Donnell, L. Mechanisms of spermiogenesis and spermiation and how they are disturbed. Spermatogenesis 4, e979623 (2015).

2. Miyata, H., Shimada, K., Kaneda, Y. & Ikawa, M. Development of functional spermatozoa in mammalian spermiogenesis. Development 151, dev202838 (2024).

3. Robinson, M., Sparanese, S., Witherspoon, L. & Flannigan, R. Human in vitro spermatogenesis as a regenerative therapy - where do we stand? Nat Rev Urol 20, 461–479 (2023).

4. Giese, A. P. J. et al. CIB2 interacts with TMC1 and TMC2 and is essential for mechanotransduction in auditory hair cells. Nat Commun 8, 43 (2017).

5. Jeong, H. et al. Structures of the TMC-1 complex illuminate mechanosensory transduction. Nature 610, 796–803 (2022).

6. Blendy, J. A., Kaestner, K. H., Weinbauer, G. F., Nieschlag, E. & Schütz, G. Severe impairment of spermatogenesis in mice lacking the CREM gene. Nature 380, 162–165 (1996).

7. Davis, F. M. et al. Male infertility in mice lacking the store-operated Ca2+ channel Orai1. Cell Calcium 59, 189–197 (2016).

8. Shambharkar, P. B. et al. TMEM203 Is a Novel Regulator of Intracellular Calcium Homeostasis and Is Required for Spermatogenesis. PLoS One 10, e0127480 (2015).

9. Lee, J. H., Kim, H., Kim, D. H. & Gye, M. C. Effects of calcium channel blockers on the spermatogenesis and gene expression in peripubertal mouse testis. Arch Androl 52, 311–318 (2006).

10. Wolski, K. M., Perrault, C., Tran-Son-Tay, R. & Cameron, D. F. Strength measurement of the Sertoli-spermatid junctional complex. J Androl 26, 354–359 (2005).

11. Coste, B. et al. Piezo1 and Piezo2 are essential components of distinct mechanically activated cation channels. Science 330, 55–60 (2010).

12. Kawashima, Y. et al. Mechanotransduction in mouse inner ear hair cells requires transmembrane channel-like genes. J Clin Invest 121, 4796–4809 (2011).

13. Hermann, B. P. et al. The Mammalian Spermatogenesis Single-Cell Transcriptome, from Spermatogonial Stem Cells to Spermatids. Cell Rep 25, 1650–1667.e8 (2018).

14. Pan, B. et al. TMC1 Forms the Pore of Mechanosensory Transduction Channels in Vertebrate Inner Ear Hair Cells. Neuron 99, 736–753.e6 (2018).

15. Kurima, K. et al. TMC1 and TMC2 Localize at the Site of Mechanotransduction in Mammalian Inner Ear Hair Cell Stereocilia. Cell Rep 12, 1606–1617 (2015).

16. Fu, S. et al. Mammalian TMC Family Proteins are Mechanically Gated Ion Channels. 2026.08.18.745354 Preprint at 10.64898/2026.08.18.745354 (2026).

17. Ebrahim, S. et al. Transmembrane channel-like 4 and 5 proteins at microvillar tips are potential ion channels and lipid scramblases. Preprint at 10.1101/2024.08.22.609173 (2024).

18. Wang, J. et al. TMC7 deficiency causes acrosome biogenesis defects and male infertility in mice. Elife 13, RP95888 (2024).

19. Lv, Z. et al. TMC7 is required for spermiogenesis and male fertility by regulating TGN-derived vesicles. International Journal of Biological Macromolecules 293, 139070 (2025).

20. Zhou, F., Li, Y., Zhang, J. & Wang, X. Diagnostic yield of exome sequencing in nonobstructive azoospermia (NOA): A systematic review and meta-analysis. PLoS One 20, e0338892 (2025).

21. Clark, A. G., Wartlick, O., Salbreux, G. & Paluch, E. K. Stresses at the Cell Surface during Animal Cell Morphogenesis. Current Biology 24, R484–R494 (2014).

22. Dai, J., Sheetz, M. P., Wan, X. & Morris, C. E. Membrane tension in swelling and shrinking molluscan neurons. J Neurosci 18, 6681–6692 (1998).

23. Desplat, A. et al. Piezo1-Pannexin1 complex couples force detection to ATP secretion in cholangiocytes. J Gen Physiol 153, e202112871 (2021).

24. Mulhall, E. M. et al. Direct observation of the conformational states of PIEZO1. Nature 620, 1117–1125 (2023).

25. Syeda, R. et al. Chemical activation of the mechanotransduction channel Piezo1. Elife 4, e07369 (2015).

26. Hogarth, C. A. et al. Processive pulses of retinoic acid propel asynchronous and continuous murine sperm production. Biol Reprod 92, 37 (2015).

27. Kirsanov, O. et al. Modeling mammalian spermatogonial differentiation and meiotic initiation in vitro. Development 149, dev200713 (2022).

28. Kirsanov, O. et al. Retinoic acid is dispensable for meiotic initiation but required for spermiogenesis in the mammalian testis. Development 150, dev201638 (2023).

29. Kierszenbaum, A. L., Rivkin, E. & Tres, L. L. Molecular biology of sperm head shaping. Soc Reprod Fertil Suppl 65, 33–43 (2007).

30. Dai, J. et al. IQCN disruption causes fertilization failure and male infertility due to manchette assembly defect. EMBO Mol Med 14, e16501 (2022).

31. Keller, T. C. & Mooseker, M. S. Ca++-calmodulin-dependent phosphorylation of myosin, and its role in brush border contraction in vitro. J Cell Biol 95, 943–959 (1982).

32. Dong, Z. et al. TRPM-mediated mechanoreception regulates myosin oscillation during tissue elongation. Curr Biol 35, 5793–5807.e4 (2025).

33. Vicente-Manzanares, M., Ma, X., Adelstein, R. S. & Horwitz, A. R. Non-muscle myosin II takes centre stage in cell adhesion and migration. Nat Rev Mol Cell Biol 10, 778–790 (2009).

34. Fettiplace, R., Furness, D. N. & Beurg, M. The conductance and organization of the TMC1-containing mechanotransducer channel complex in auditory hair cells. Proc Natl Acad Sci U S A 119, e2210849119 (2022).

35. Li, Y. et al. Structural insights into calcium-dependent CIB2-TMC1 interaction in hair cell mechanotransduction. Commun Biol 8, 306 (2025).

36. Zheng, W. & Holt, J. R. The Mechanosensory Transduction Machinery in Inner Ear Hair Cells. Annu Rev Biophys 50, 31–51 (2021).

37. Yuan, W. et al. CIB1 Is Essential for Mouse Spermatogenesis. Mol Cell Biol 26, 8507– 8514 (2006).

38. Xu, Z. et al. CIB4 is essential for the haploid phase of spermatogenesis in mice†. Biol Reprod 103, 235–243 (2020).

39. Ebrahim, S. et al. Stereocilia-staircase spacing is influenced by myosin III motors and their cargos espin-1 and espin-like. Nat Commun 7, 10833 (2016).

40. Labay, V., Weichert, R. M., Makishima, T. & Griffith, A. J. TOPOLOGY OF TRANSMEMBRANE CHANNEL-LIKE GENE 1 PROTEIN (TMC1). Biochemistry 49, 8592–8598 (2010).

41. Soler, D. C. et al. An uncharacterized region within the N-terminus of mouse TMC1 precludes trafficking to plasma membrane in a heterologous cell line. Sci Rep 9, 15263 (2019).

42. Abramson, J. et al. Accurate structure prediction of biomolecular interactions with AlphaFold 3. Nature 630, 493–500 (2024).

43. Mayr, B. & Montminy, M. Transcriptional regulation by the phosphorylation-dependent factor CREB. Nat Rev Mol Cell Biol 2, 599–609 (2001).

44. Martianov, I. et al. Cell-specific occupancy of an extended repertoire of CREM and CREB binding loci in male germ cells. BMC Genomics 11, 530 (2010).

45. Kosir, R. et al. Novel Insights into the Downstream Pathways and Targets Controlled by Transcription Factors CREM in the Testis. PLoS One 7, e31798 (2012).

46. Murthy, S. E. et al. OSCA/TMEM63 are an Evolutionarily Conserved Family of Mechanically Activated Ion Channels. Elife 7, e41844 (2018).

47. Mundt, N., Spehr, M. & Lishko, P. V. TRPV4 is the temperature-sensitive ion channel of human sperm. eLife 7, e35853 (2018).

48. Platts, A. E. et al. Success and failure in human spermatogenesis as revealed by teratozoospermic RNAs. Hum Mol Genet 16, 763–773 (2007).

49. Kui, F., Ye, H., Chen, X.-L. & Zhang, J. Microarray meta-analysis identifies candidate genes for human spermatogenic arrest. Andrologia 51, e13301 (2019).

50. Pacheco, S. E. et al. Integrative DNA methylation and gene expression analyses identify DNA packaging and epigenetic regulatory genes associated with low motility sperm. PLoS One 6, e20280 (2011).

51. Wagner, E. L. et al. Repair of noise-induced damage to stereocilia F-actin cores is facilitated by XIRP2 and its novel mechanosensor domain. eLife 12, e72681 (2023).

52. Kim, C. R., Noda, T., Okada, Y., Ikawa, M. & Baek, S. H. Protocol for isolation of spermatids from mouse testes. STAR Protocols 2, 100254 (2021).

53. Velte, E. K. et al. Differential RA responsiveness directs formation of functionally distinct spermatogonial populations at the initiation of spermatogenesis in the mouse. Development 146, dev173088 (2019).

54. R Core Team. R: A Language and Environment for Statistical Computing. R Foundation for Statistical Computing (2024).

55. Hao, Y. et al. Dictionary learning for integrative, multimodal and scalable single-cell analysis. Nat Biotechnol 42, 293–304 (2024).

56. Young, M. D. & Behjati, S. SoupX removes ambient RNA contamination from droplet-based single-cell RNA sequencing data. Gigascience 9, giaa151 (2020).

57. Korsunsky, I. et al. Fast, sensitive and accurate integration of single-cell data with Harmony. Nat Methods 16, 1289–1296 (2019).

58. Cao, J. et al. The single-cell transcriptional landscape of mammalian organogenesis. Nature 566, 496–502 (2019).

59. Qiu, X. et al. Reversed graph embedding resolves complex single-cell trajectories. Nat Methods 14, 979–982 (2017).

60. Trapnell, C. et al. The dynamics and regulators of cell fate decisions are revealed by pseudotemporal ordering of single cells. Nat Biotechnol 32, 381–386 (2014).

61. Butler, A., Hoffman, P., Satija, R. & Stuart, T. SeuratWrappers. GitHub (2024).

62. Wickham, H. et al. Welcome to the Tidyverse. Journal of Open Source Software 4, 1686 (2019).

63. Wickham, H. Ggplot2: Elegant Graphics for Data Analysis. (Springer, Cham, 2016). doi:10.1007/978-3-319-24277-4.

64. Blighe, K., Rana, S. & Lewis, M. EnhancedVolcano: publication-ready volcano plots with enhanced colouring and labeling. Bioconductor (2024).

65. Insights into substrate binding and utilization by hyaluronan synthase. eLife 14, RP109624 (2026).

66. Lehe, M. D., Almofeez, R., Jeffery, E. D. & Sheynkman, G. M. Advances in Mass Spectrometry Instrumentation and Methodology for Analysis of Alternative Protein Isoforms. J Mass Spectrom 61, e70024 (2026).

67. Naydenov, N. G., et al. The septin cytoskeleton is a regulator of intestinal epithelial barrier integrity and mucosal inflammation. JCI Insight 10, e191538 (2025).

68. Meistrich, M. L. & Hess, R. A. Assessment of spermatogenesis through staging of seminiferous tubules. Methods Mol Biol 927, 299–307 (2013).

69. Niedenberger, B. A. & Geyer, C. B. Advanced immunostaining approaches to study early male germ cell development. Stem Cell Res 27, 162–168 (2018).

